# Structural and molecular basis for foot-and-mouth disease virus neutralization by two potent protective antibodies

**DOI:** 10.1101/2020.12.31.424923

**Authors:** Hu Dong, Pan Liu, Manyuan Bai, Kang Wang, Rui Feng, Dandan Zhu, Yao Sun, Suyu Mu, Haozhou Li, Michiel Harmsen, Shiqi Sun, Xiangxi Wang, Huichen Guo

## Abstract

Outbreaks of Foot-and-mouth disease (FMD) caused by FMD virus result in significant economic losses. Vaccination is helpful, but the benefits are diminished with antigenic diversity within serotypes, instability of the immunogen and inability to confer protection for long durations. Here we have further dissected the mechanisms underpinning the protective efficacy of two previously reported neutralizing antibodies (NAbs), M8 and M170. The atomic details of the epitopes of M8 and M170 unveiled suggest that protection is conferred by disrupting the virus-receptor interactions. Consequently, administration of these NAbs conferred prophylactic and therapeutic benefit in guinea pigs, raising the possibility of administering NAbs before or during vaccination to confer immediate protection; well before the bolstering of the immune response by the vaccine. Differences in the residues and the conformation of elements making up the epitopes explain the differences in specificities of M8 and M170. An ability to bind 146S viral particles specifically, but not 12S degraded components, highlights a likely role for M170 in the quality control of vaccines.

## Introduction

Foot-and-mouth disease (FMD) is an economically devastating and highly contagious viral disease of cloven-hoofed animals with a global distribution. The causative agent, FMD virus (FMDV) is a small non-enveloped RNA virus, belonging to the *Aphthoviruses* genus within *Picornaviridae* family(Tuthill et al., 2010). Seven immunologically distinct serotypes (O, A, C, Asia 1, SAT1, SAT2 and SAT3) of FMDV and many variants with a spectrum of antigenic diversity have been identified(Knowles and Samuel, 2003). Among these, serotype O poses the most serious global threats. Control of FMD has been largely reliant on vaccinations with inactivated virus vaccines. However, significant antigenic diversity within FMDV serotypes and inability of the vaccines to induce immune protection for a long duration of time impinge on the efficacy of available vaccines. The roles of neutralizing antibodies (NAbs) as the principal protective components of the immune responses to FMDV vaccination or infection have been well established (Juleff et al., 2009; Pay and Hingley, 1987). Passive immunization of NAbs has also been demonstrated to be effective in curing FMD and many viral diseases (Hangping Yao, 2020; Harmsen et al., 2008; Harmsen et al., 2007; Li Zhang, 2020; Ling Zhu, 2020; Lv et al., 2020; Nan Wang, 2020; Qiu et al., 2018). A deep understanding of the molecular basis for viral neutralization by antibodies and the identification of key viral epitopes would aid in the development of potent rationally designed broad-spectrum vaccine.

FMDV possesses an icosahedral capsid composed of 60 copies each of four structural proteins (VP1-VP4), among which VP1-VP3 constitute the outer shell, while VP4 lines the interior surface (Acharya et al., 1989; Fry et al., 2005). Structures of several serotypes of FMDV reported previously reveal that although the overall structure is conserved between the serotypes, the surface-exposed loops joining the strands of the jelly rolls differ substantially between serotypes. Immunological epitopes have been mostly found to be located on these loops (Hewat et al., 1997). Numerous studies have been carried out in the past two decades for mapping antigenic sites on the viral surface *via* NAb-resistant mutants (Asfor et al., 2014; Ciurea et al., 2000; Crowther et al., 1993; Kitson et al., 1990; Opperman et al., 2014). For FMDV type O, at least five neutralizing antigenic sites have been revealed, of which VP1 GH, an exceptionally long flexible loop, not only makes up a large portion of a major antigenic site, but also contains a conserved RGD (Arg-Gly-Asp) motif, essential for the attachment of the virus to the cellular receptor integrin (Berryman et al., 2005; Burman et al., 2006; Logan et al., 1993). Understanding of the mechanisms by which NAbs neutralize FMDV and the “escape” of the viral variants from neutralization by the NAb requires a detailed analysis of virus-antibody interactions at atomic resolution. Although cryo-electron microscopy (cryo-EM) structures of FMDV-NAb complexes at low resolution (~30 Å) and crystal structures of viral peptide-antibody complexes have been reported (Hewat et al., 1997; Verdaguer et al., 1995; Verdaguer et al., 1999), these structures neither provide “smoking gun” evidence to clarify the mechanism of neutralization nor do they precisely define the atomic determinants of the interaction. Currently, there is no high-resolution structural information available for FMDV in complex with NAbs.

Here we revisited two “old” llama single-domain antibodies, M8 and M170, reported 15 years ago (Harmsen et al., 2007), in order to study the molecular mechanisms of neutralization of FMDV by these antibodies at the atomic level. Both NAbs can neutralize FMDV type O infection at sub-*μM* level by blocking viral attachment to the host cell and interfering with viral uncoating. Atomic structures of FMDV in complex with M8/M170 reveal the nature of the binding modes, location and conformation of epitopes targeted by these two antibodies. These *bona fide* epitopes, together with the results of the immunological studies, facilitate the identification of the areas of antigenic variability and conservation among serotypes of FMDV The atomic details of immunogens unveiled in this study can inform strategies for rationally designing effective broad-spectrum vaccines and antiviral therapeutics against FMD.

## Results

### Characterizations of anti-FMDV NAbs M8 and M170

By using phage display immune libraries derived from four llamas, we previously identified 24 single-domain antibodies capable of neutralizing FMDV type O *in vitro* (Harmsen et al., 2007). Among these, M8 and M170 exhibited relatively strong neutralizing activities (Harmsen et al., 2007). However molecular details of mechanisms of neutralization of FMDV by these antibodies are unclear. Here, we systematically analyzed the immunological properties of these two NAbs and delineated the molecular basis of neutralization against FMDV infection using a combination of immunological and virological assays, structural analysis, biochemical and animal studies. To investigate the serotype specificity or cross-reactivity of M8 and M170, we propagated and purified FMDV O, A, Asia 1 as well as C serotype strains and separately examined their binding abilities to each antibody by enzyme-linked immunosorbent assay (ELISA). The ELISA results revealed that M170 only binds to type O, but M8 is capable of reacting with all the indicated serotypes, suggesting that M8 and M170 are type cross-reactive and O-specific, respectively (Fig. 1a). Surface plasmon resonance (SPR) experiments verified that both M8 and M170 interact with type O with a high binding affinity of 0.5 and 1.0 nM, respectively (Fig. 1b). To explore whether these two antibodies can simultaneously bind the virus, we performed a competitive SPR assay and the results indicated that the binding of one antibody blocks the attachment of the other (Fig. 1c), which suggests that M8 and M170 compete with each other for simultaneous binding albeit with different characteristics in binding distinct serotypes and targeting distinct antigenic sites. Cell-based neutralization assays showed that both M8 and M170 exhibit potent neutralizing activities against type O with a 50% neutralizing concentration value (Neut50) of 0.8 and 3.2 μM, respectively (Fig. 1d). Perhaps correlated with the inability of simultaneous binding, the cocktail of M8 and M170 did not exhibit synergistic neutralization activity (Fig. 1e). Interestingly, results of a fluorescence-based assay revealed that M8, rather than M170, destabilized FMDV particles by 3-8 °C in an incubation time dependent manner at physiological conditions (pH 7.5), indicating that physical destabilization of viral particles that interferes with normal uncoating may be a possible neutralization mechanism for M8 (Fig. 1f). Such destabilization of viral particles has been also observed in studies dissecting mechanisms of NAbs neutralizing human enterovirus A71 (EV71) and other picornaviruses (Plevka et al., 2014; Wang et al., 2017; Zhu et al., 2018b).

**Fig. 1.**
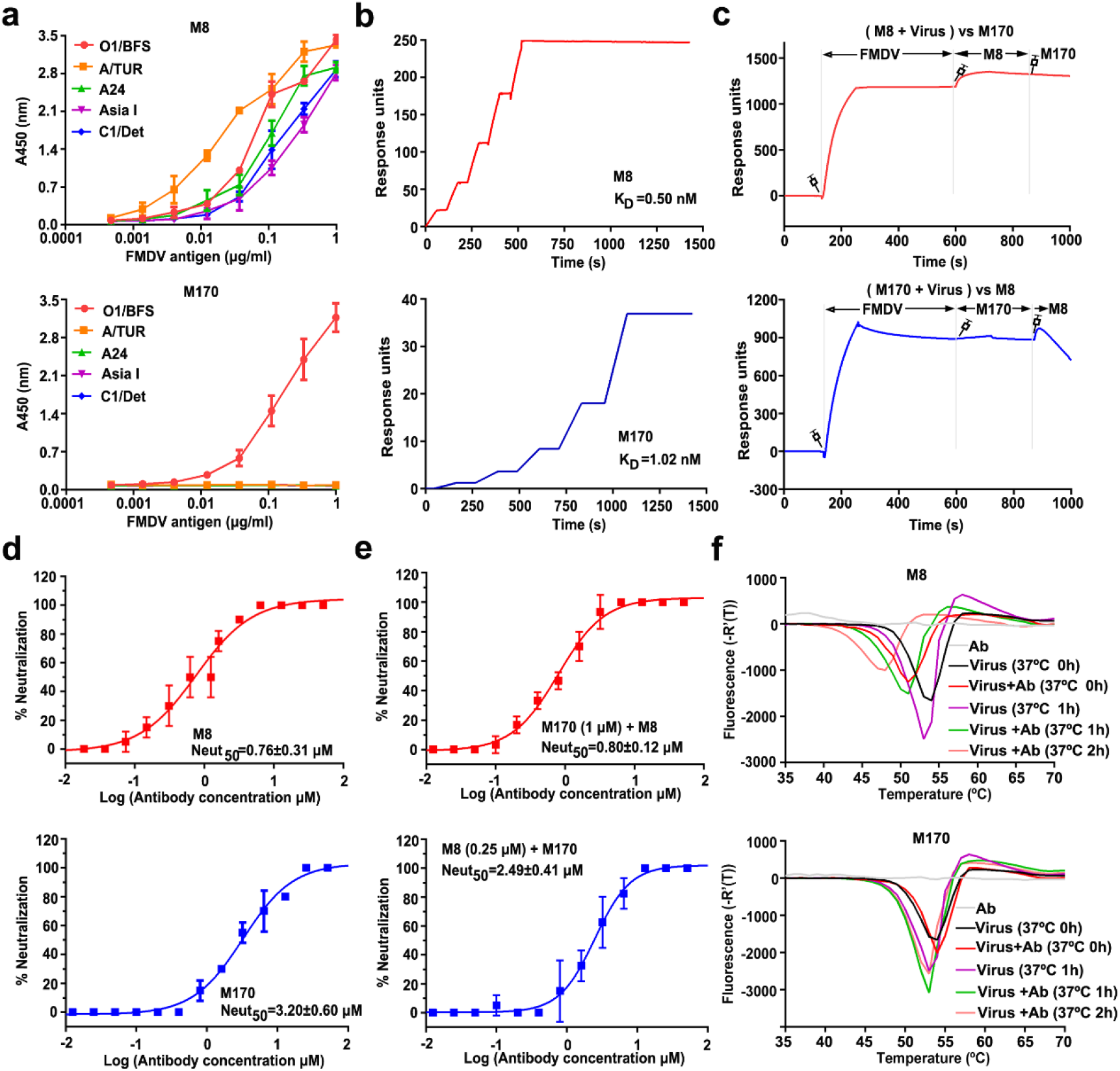
Characterization of M8 and M170. **a** Dose-dependent binding analysis of M8 and M170 against representative FMDV serotypes by ELISA. Each plot represents the mean of OD450 values from triplicate wells. Error bars represent mean ± SD. FMDV O (O1/BFS1860/UK/67), A (A/TUR/20/2006, A24/Cruzeiro/BRA/55), Asia 1 (Asia 1/Shamir/ISR/89) and C (C1/Detmold/FRG/60). **b** BIAcore SPR kinetic profile of mAb M8 (top) and M170 (bottom) against FMDV O. The binding affinity is depicted by KD (equilibrium dissociation constant, K_D_ = kd/ka), which was calculated by the BIAcore 3000 analysis software (BIAevaluation version 4.1). **c** Analysis of the simultaneous binding of M8 and M170 to FMDV through a competitive SPR assay. Due to the requirement of acidic condition for the sensor-labeling, M8 or M170, rather than FMDV, was loaded onto the sensor. In the upper panel, the sensor was labeled with M8. FMDV was inject first, followed by the second injection of M8 again to fully occupy all binding sites for M8 on viral surface, then M170 flowed through. In the bottom panel, the sensor was labeled with M170. The SPR procedure of M170 is same to that of M8. Note: signals (during 800-1,000s) decreased upon M8 binding in the bottom panel, which might result from the ability that M8 has the potential to strip FMDV off the M170-labelled sensor due to the higher binding affinity. **d** Neutralization of FMDV O by M8 (top) and M170 (bottom) using plaque-reduction neutralization assay. Neut_50_ values indicate concentration of antibody required to neutralize fifty percent of the viral titer. The Neut50 of M8 and M170 were 0.8 μM and 3.2 μM, respectively. **e** Neutralizing activities of the combination of M8 and M170. The Data is presented as the mean ± SD of triplicate measurements. **f** Stabilities of FMDV O upon addition of M8 (top) or M170 (bottom) at physiological temperature. The release of RNA reflecting the dissociation of capsids was detected by an increase in the fluorescence signal (SYTO9 fluorescent).

### Prophylactic and therapeutic efficacy of M8 and M170 in guinea pigs

Given the potent neutralizing activities of M8 and M170 at sub-μM concentrations, we next sought to assess the *in vivo* protection efficacy of these two antibodies in animal model. Guinea pigs (GP) susceptible to experimental FMDV infections have been widely used as preferential laboratory species for FMDV research since the clinical outcome resembles that observed in natural hosts when compared to other animal models (Brown, 2003; Tekerlekov et al., 1976). Inoculation of FMDV type O (100 TCID50) in the hind footpad led to productive replication of the virus and developed generalized FMD symptoms (lesions at the inoculation sites) at 3 day post infection (PI). Guinea pigs were administered M8 or M170 1 day before or after inoculation of the FMDV, representing a prophylactic (P) or therapeutic (T) setting, respectively (Fig. 2a). In the prophylactic mode, a single intramuscular injection of M8 at 2.5 mg/kg protected 75% of the animals from FMD until the end of the experiments and delayed morbidity in rest of the animals exhibiting symptoms of FMD when compared to the control. In contrast to this, only 25% of the animals receiving the same dose of M170 were protected from FMDV infections, while 75% of the animals developed FMD symptoms later at day 4 PI (Fig. 2b). In line with these results, viral loads in the blood were nearly undetectable during the course of the monitoring of the infection in M8 administrated group. However, a severe viremia occurred at day 2 PI in control group. Pre-treatment of M170 alleviated the viremia and cleared ~95% of the viral titer when compared to the control group at day 3 PI (Fig. 2c). Therapeutic administration of either M8 or M170 one day after virus challenge resulted in a relatively weak protective efficacy with 25% protection for M8 and delayed morbidity for M170 when compared to the control (Fig. 2d). Correlated with this, high levels of viral replication prior to therapeutic administration of the NAbs were detected, whereas M8 and M170 treatment at all indicated time-points resulted in largely reduced viral titers with a clear trend for greater reduction with earlier treatment (Fig. 2e). Of note, efficient viral infection that had been established at day 1 PI may require a higher dose of antibodies for rapid and complete clearance of FMDV or an earlier therapeutic treatment for suppressing massive replication of viruses.

**Fig. 2.**
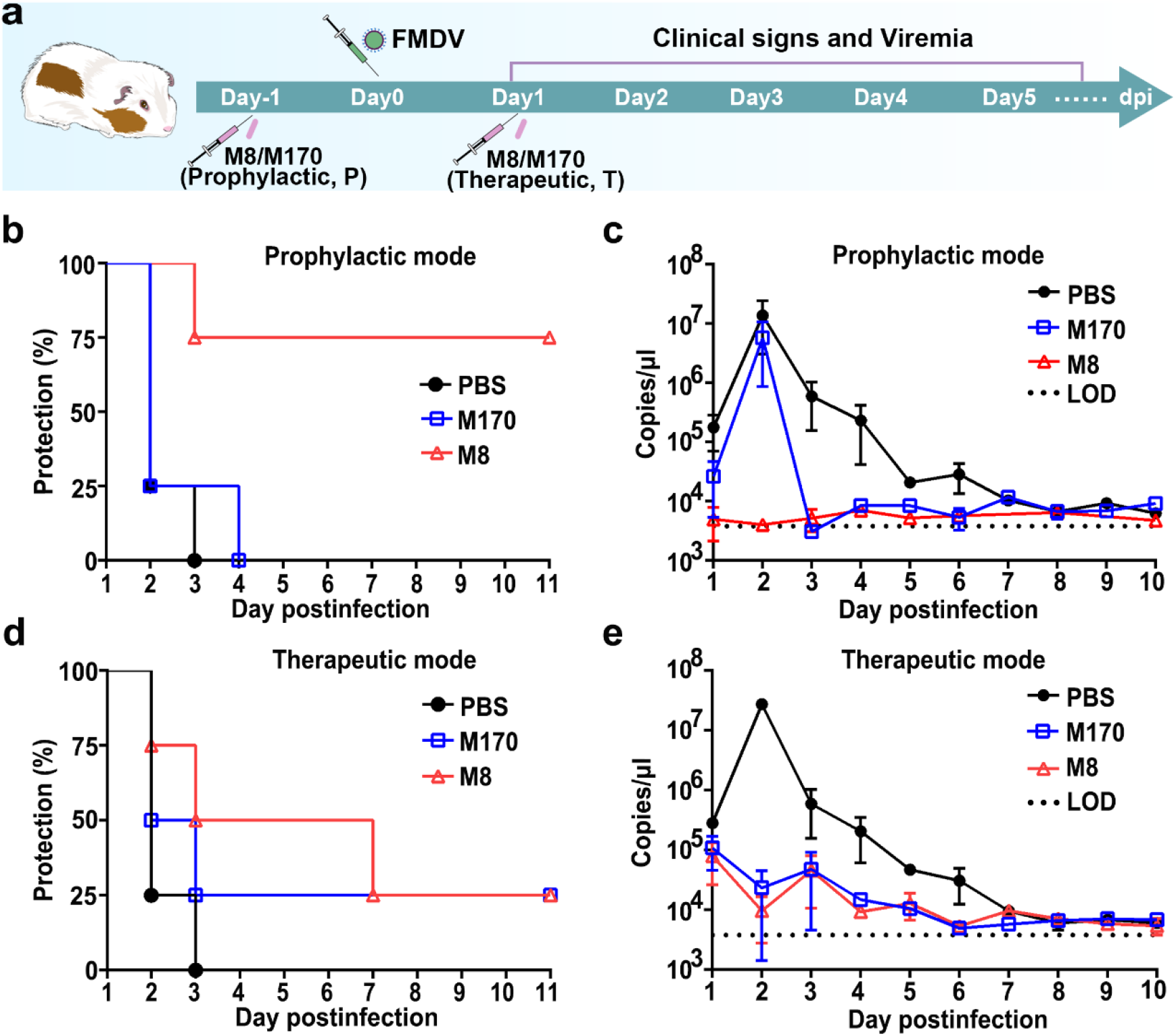
Prophylactic and therapeutic efficacy of M8 and M170. **a** Schematic diagram of M8 or M170 treatment in guinea pig model. Groups of guinea pigs (n=4) were administrated intramuscularly with M8/M170 (2.5 mg/kg) 1 day before (prophylactic) or after (therapeutic) challenge with 100 ID_50_ of FMDV on the left hind footpad. Guinea pigs injected intramuscularly with PBS before or after challenge were acted as control groups. The day of FMDV infection was marked as day 0. Protection of guinea pigs against FMDV O by passive immunization with M8 or M170 was analyzed in the prophylactic (**b**) and therapeutic (**d**) modes. No lesions on rear feet were considered as full protection. The copies of virus mRNA in the blood sample from guinea pigs of the prophylactic (**c**) and therapeutic (**e**) groups were quantified by the real-time quantitative PCR (RT-qPCR), the limit of detection (LOD) of viral mRNA in blood was labeled.

### M8 and M170 prevent viral attachment to host cells by blocking virus-receptor interaction

Integrins, heterodimeric adhesive glycosylated membrane proteins located on the surface of most cells, have been identified as FMDV receptors that bind viruses via a small cleft at the subunit interface of the head (Kotecha et al., 2017). FMDV enters host cells by integrin-mediated endocytosis, then disassociates into pentamers to release RNA in the acidic environment of the endocytic compartments, differing from uncoating intermediates observed in many other picornaviruses (Bostina et al., 2011; Wang et al., 2012; Zhu et al., 2018a). To investigate whether M8 or M170 interferes with the binding of FMDV to integrin, we performed competitive SPR-based binding assays. We chose to load integrin onto the CM5 sensor due to the requirement of acidic condition for the sensor-labeling and washed the sensor with neutral buffer solutions. Mixtures of FMDV particles with various concentrations of M8 or M170 acted as running phase to flow through the sensor. The results indicated that both M8 and M170 can prevent FMDV from binding integrin in a concentration-dependent manner (Fig. 3a). Besides, the blocking of binding by M8 appeared more potent when compared to M170, which explains the differential neutralizing activities of these two antibodies (Fig. 3a). To further verify these results in cell-based viral infection model, we used real-time reverse transcriptase-PCR (RT-PCR) to quantify the amount of virus remaining on the surface of cells that were treated with M8 or M170 pre- and post-viral attachment at 4 °C. Consistent with the competitive binding assay results, both M8 and M170 efficiently prevented FMDV type O attachment to the cell surface and could displace the viral particles that had already bound to the cell surface in a dose-dependent manner (Fig. 3b and 3c).

**Fig. 3.**
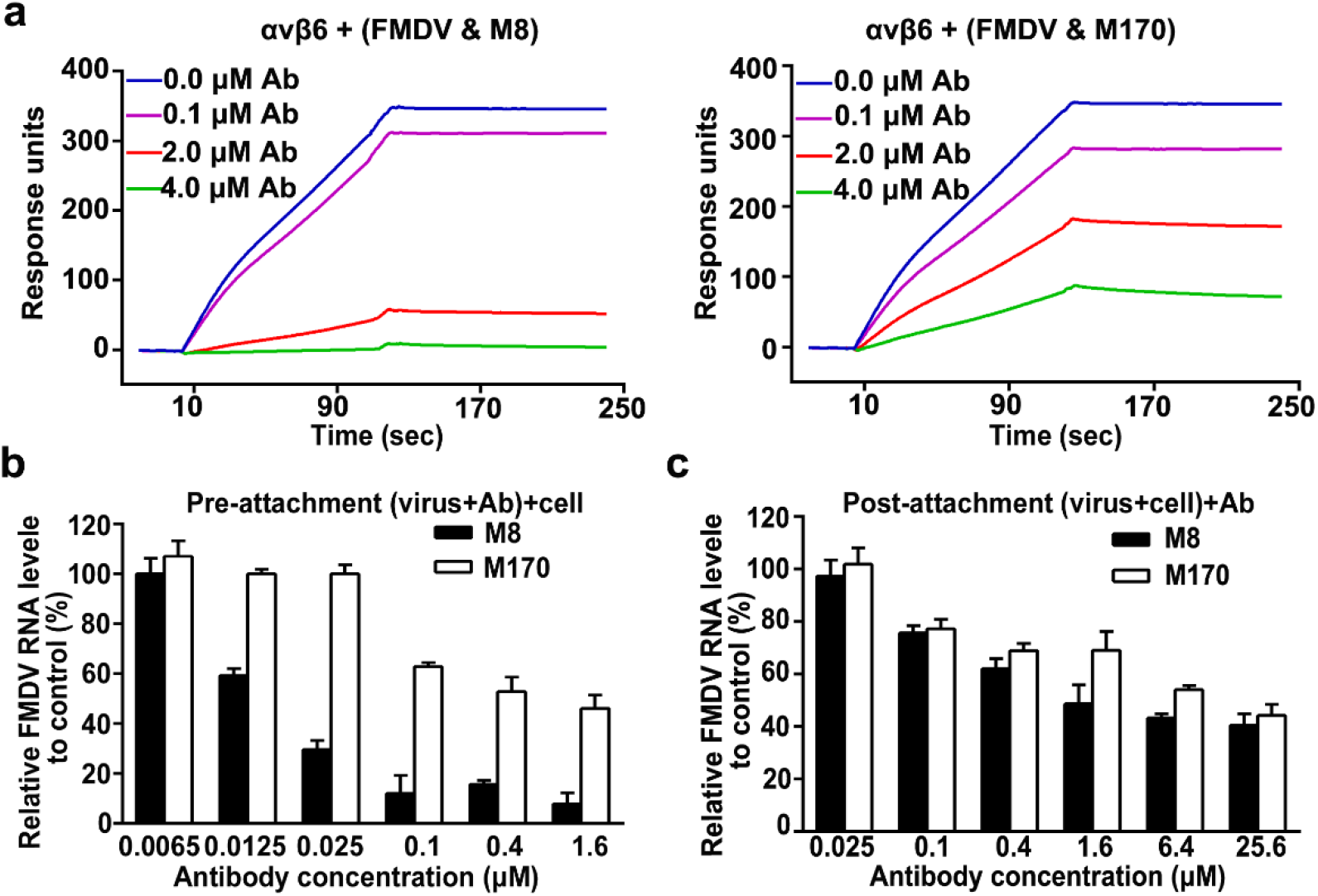
M8 and M170 block the interactions of FMDV with integrin. **a** BIAcore SPR kinetics shows the competitive binding of M8/M170 and αvβ6 integrin to FMDV O. For both panels, αvβ6 integrin was immobilized onto the sensor chips. Mixtures of FMDV O (20 nM) with various concentrations of M8 (left) or M170 (right) acted as running phase to flow through the sensor. Binding signals were detected. **b** Amount of virions remaining on the cell surface, as detected by real-time PCR, when exposed to M8 or M170 before or after the virions attach to BHK21 cells. Data is presented as the mean ± SD. Experiments were repeated in triplicate.

### Structures of FMDV in complex with its NAbs M8 and M170

To define the key epitopes and atomic determinants of the interactions between FMDV and its two NAbs precisely, we determined the cryo-EM structures of FMDV type O in complex with M8 or M170. Cryo-EM micrographs of the purified formaldehyde-inactivated FMDV mature particles in complex with M8 or M170 were recorded using an FEI Titan Krios electron microscope equipped with a Gatan K2 Summit detector (Supplementary Fig. 1). Icosahedral reconstructions of FMDV-M8 and FMDV-M170 complexes were achieved at resolutions of 3.2 Å and 3.1 Å, respectively (Supplementary Fig. 2). The maps for the viral capsid were of sufficient resolution to build and refine the atomic model. But, the densities for antibodies and binding interface were relatively weak due to the unsaturated occupancy and conformational heterogeneity (Supplementary Fig. 3), which is in line with the structural observations of flexible bindings of integrins to FMDV at different angles (Kotecha et al., 2017). To improve the resolution of the binding interface, local refinement by using an optimized “block-based” reconstruction approach (Wang et al., 2019; Yang et al., 2020; Yuan et al., 2018) was performed. This led to an improvement of the local resolution to 3.9 Å and 3.5 Å for the binding interfaces of FMDV-M8 and FMDV-M170, respectively, enabling a reliable analysis of the interaction modes (Supplementary Fig. 4).

As expected, M8 and M170 recognize distinct antigenic sites (Fig. 4). M8 binds to the FMDV viral surface within the pentameric building blocks surrounding the fivefold axis at a position akin to the “canyon” in enteroviruses (Dang et al., 2014) (Fig. 4a–4d). The location of this binding site is similar to those observed previously for 11G1 antibody bound to Enterovirus D68 (EVD68) and NAb 4B10 bound to Echovirus 30 (E30) (Kang Wang, 2020a; Zheng et al., 2019). In contrast, M170 targets the surface along the edges of pentameric building block of the virus (Fig. 4a–4d), nearby the twofold axis at a position similar to that observed for D6 antibody bound to EV71(Zhu et al., 2018b). Apart from targeting distinct antigenic sites, M8 and M170 adopt significantly different configurations upon binding to the FMDV (Fig. 4e). When viewed down the fivefold axis, individual M8 stands vertically, mimicking the association mode between the β subunit of integrin and FMDV (Kotecha et al., 2017), while M170 locates aside M8, adjacent to the twofold axis, but inclines by ~45° towards the viral surface in an attachment mode resembling the strategy used by the α subunit of integrin for its interaction with FMDV (Fig. 4e). However, minor steric clashes resulting from one overlapped residue within the binding sites and the inclined posture of M170 prevent the simultaneous binding of M8 and M170 to FMDV (Fig. 4f), thereby failing to make up a non-competing pair of antibody cocktail.

**Fig. 4.**
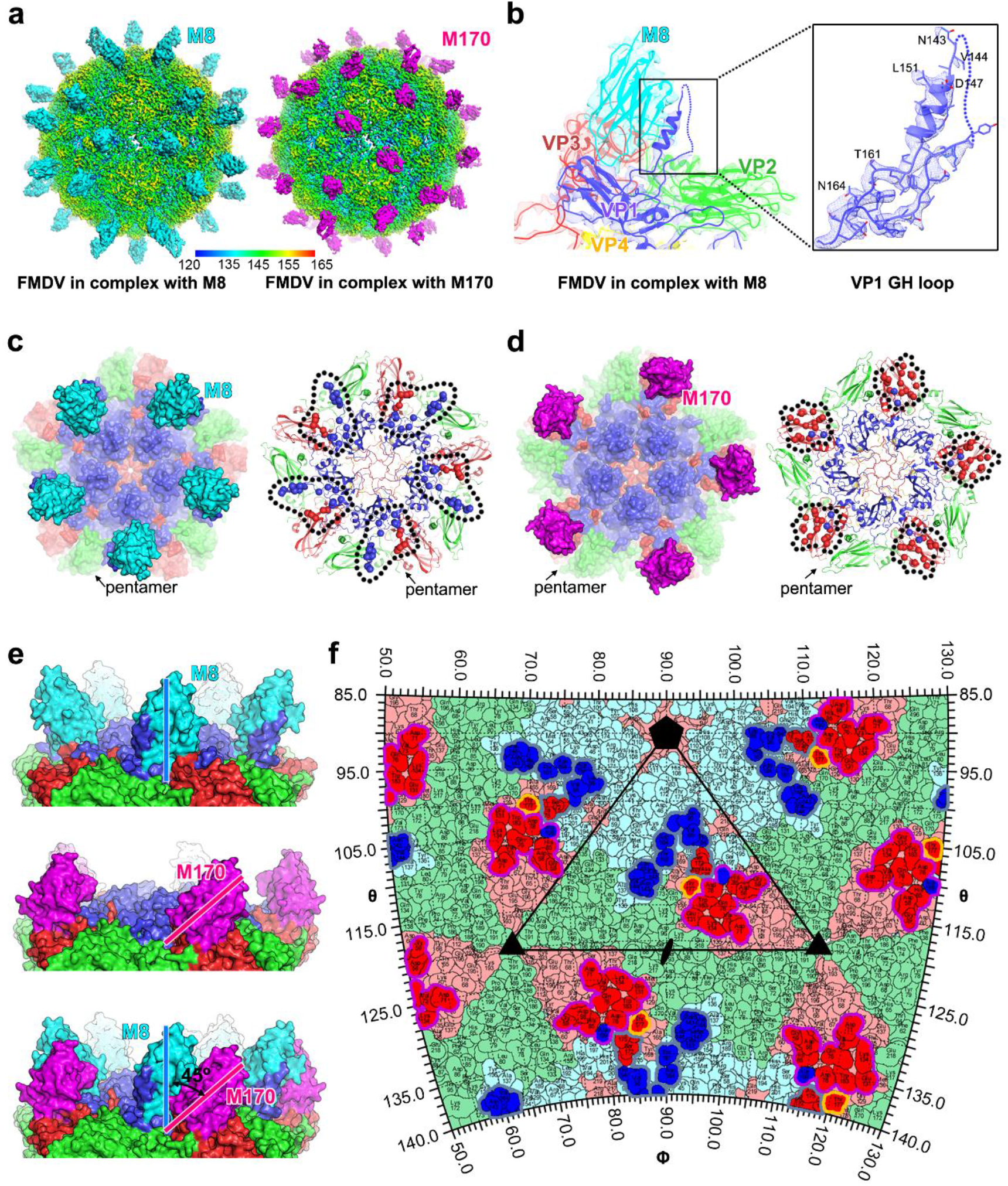
Cryo-EM structures of FMDV in complex with M8 or M170. **a** Cryo-EM maps of FMDV-M8-complex (left) and FMDV-M170-complex (right). The viral capsids of both complexes are rainbow-colored as the color bar shown below; the M8 and M170 are colored in cyan and magenta, respectively. **b** Electron density maps for the binding interface of M8 and one protomer of the FMDV capsid. The inset shows the density maps and atomic model of VP1 GH loop. Residues with side chains are labeled. **c** and **d** Surface representation (left) and epitopes (right) of M8 (c) and M170 (d) on a viral pentamer. The pentamers are shown as surface (left) or cartoon (right) in the signature colors (VP1, blue; VP2, green; VP3, red), while M8 and M170 are colored in the same scheme as in Fig. 4a. The epitopes of M8 (right in 4c) and M170 (right in 4d) are shown as spheres and those from one protomeric unit are circled by black dotted lines. **e** Side views of two NAbs bound to a pentamer, up: FMDV-M8 complex, and middle: FMDV-M170 complex, down: the superimposition of two complexes shows an angle of ~45° between M8 and M170. The same color scheme is applied as above. **f** The M8 and M170 footprints on the FMDV surface. Residues of VP1, VP2, and VP3 are outlined in blue, green, and red, respectively. Residues involved in binding to NAbs are shown in brighter colors corresponding to the protein chain they belong to, the footprints of M8 and M170 are indicated by gray and magenta lines, respectively. The overlapped residue for binding to both M8 and M170 is marked by yellow lines.

### Epitopes of NAbs M8 and M170

Analysis of the structures of FMDV in complex with M8 or M170 helped identify the epitopes of these two antibodies. M8 recognizes a conformational epitope constructed by VP1 EF, VP1 GH and VP3 GH loops within a protomer (Fig. 5a and Supplementary Fig. 5), which differs from previously reported epitopes of an antibody escape mutant where the VP1 GH loop functions as either an independent or as a discontinuous epitope encompassing the C terminus of VP1 (Crowther et al., 1993). All three complementary determining regions (CDR): CDR1 (residues 32-39), CDR2 (residues 60) and CDR3 (residues 110-119) and framework region (FR, residues 66-70) of M8 are involved in extensive binding contacts with VP1 and VP3, burying ~1,100 Å^2^ of surface area by pinching the helix of VP1 GH loop in the “up” conformation between the CDR2 and CDR3 (Fig. 4b, Fig. 5b and Supplementary Fig. 6). Notably, VP1 GH loop, highly disordered in the native virus structures, adopts a more ordered position lying along the viral surface when the disulfide bond linking the base of the loop (C134) to C130 of VP2 is reduced (Logan et al., 1993). The native high-resolution structural details of this loop enable us to define precisely the atomic determinants of the virus-antibody interaction. The epitope recognized by M8 primarily locates in the VP1 EF loop (A93, P94, V95 and L98), VP1 GH loop (K133, N143, V144, D147, L148, Q149, L151, A152, Q153, T161, N164 and A167), and the VP3 GH loop (A171, S172, A175, E176 and T177) (Fig. 5b). When compared to M8, M170 primarily binds to VP3 via recognizing the “knob”, βC, BC, CD, EF and GH loops of VP3, covering most of the exposed surface of VP3, and VP1 C-terminus (Fig. 5a and Supplementary Fig. 5), which is way far beyond the previously reported antigenic site 4 within VP3 (McCahon et al., 1989). The M170 paratope is composed of CDR1 (residues 30-35), CDR2 (residues 53-56) and CDR3 (residues 103-110), which bury a surface area of about ~900 Å^2^ at the interface with VP3 and VP1 through hydrophilic interactions and hydrophobic contacts (Fig. 5a and 5b). The epitope of M170 mainly includes residues 57-59 of the βB “knob”, residues 71 and 73 of BC loop, residue 76 of βC, residues 78 and 85 of CD loop, residues 131 and 134 of EF loop, residues 177, 178 and 183 of GH loop in VP3 and residues 199-202 of VP1 C-terminus (Fig. 5b). Tight binding between the antibodies and FMDV is accomplished by forming extensive hydrophilic interactions, including hydrogen bonds and salt bridges (Supplementary Table 2 and Table 3).

**Fig. 5.**
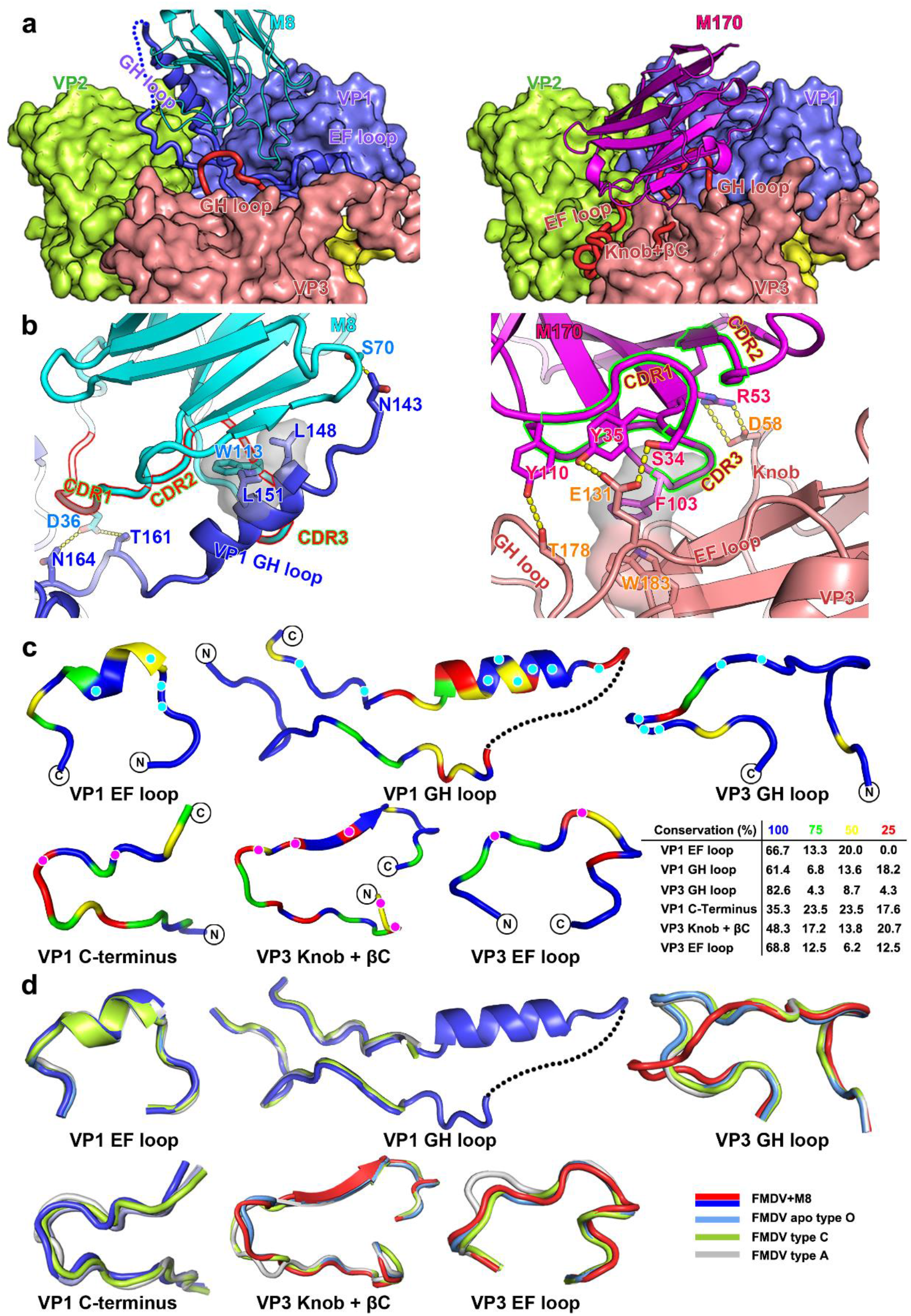
Interactions between FMDV and M8/M170. **a** The structural basis of M8 (left) and M170 (right) binding to FMDV. Loops from FMDV involved in interactions with M8/M170 are shown as cartoon, while the remaining parts are shown as surface. M8 and M170 are represented as cartoon. The color scheme is as in Fig. 4a. **b** Detailed interactions between FMDV and M8 (left), as well as FMDV and M170 (right). Residues involved in the binding are shown as sticks and hydrogen bonds are shown as yellow dashed lines. The complementary determining regions of M8 and M170 involved in the interaction with viral capsids are outlined in red and green, respectively. **c** Sequence conservation analysis. The loops from FMDV O that are involved in interactions between FMDV and M8 (VP1 GH loop, VP1 EF loop, VP3 GH loop), as well as FMDV and M170 (VP1 C-terminus, VP3 Knob-βC and VP3 EF loop) are aligned with those of 3 other FMDV serotypes (A, Asia 1 and C) and colored according to sequence conservation as listed in the table below. Residues involved in binding to M8 and M170 are marked by cyan and magenta dots. **d** Structural conservation analysis. The same loops in 4c colored in the signature color scheme are superposed with their counterparts from the *apo* O (sky blue), A (gray) and C (lime).

### M8 and M170 differ in their modes of binding and neutralization

Many NAbs with potent neutralizing activities are virus specific. Cross-reactive NAbs, in general, exhibit relatively weak antiviral activities. By contrast, in our case, M8, capable of cross-binding to several FMDV serotypes, including O, A, Asia 1 and C strains, shows better neutralizing activity than M170 that is a type O-specific antibody (Fig. 1). More importantly, cross-reactive NAbs, such as M8, primarily targeting more conserved epitopes offer clues for identifying the Achilles’ heel of viruses between serotypes, even between genera, since these regions can be targeted for the development of antiviral drugs or broad-spectrum vaccine design. To decipher the structural basis for serotype-specific and cross-reactive binding of M8 and M170, sequence and structural alignments of these four FMDV serotypes focusing on key epitopes were performed. As a major antigenic site, VP1 GH loop bears, to some extent, sequence and structural diversity within FMDV serotypes. Therefore, VP1 GH loop together with other antigenic sites can be used to distinguish type O from other serotypes. However, this region also contains a conformationally conserved helix (disordered in the native structures) with a conserved RGD motif within serotypes for receptor binding (Fig. 5c and 5d). Notably, ~70% of the binding interface between M8 and VP1 GH loop is made up by conserved residues among serotypes. Incidentally, VP1 EF and VP3 GH loops exhibit even higher conservation both in sequence and configuration (Fig. 5c and 5d). Interestingly, VP3 GH, rather than other loops, adopts an alternative configuration upon M8 binding, whilst the binding of M170 does not mediate any notable conformational change although the NAb does target VP3 GH loop as well (Supplementary Fig. 7). Overall, out of the 21 residues in the M8 epitope, 15 (~72 %) are identical among these four serotypes (Supplementary Fig. 5), which explains the cross-reactivity and comparable binding affinities. In contrast to M8, M170 buries ~680 Å^2^ of the VP3 surface by interaction with the “knob”, BC, βC, CD, EF and GH loops as well as ~220 Å^2^ of VP1 via association with its C-terminal loop. Except for VP3 GH loop, subtle conformational variations in the binding areas for M170 are observed within these four serotypes and most residues necessary for M170 binding are not conserved (Fig. 5c and 5d). The specificity of the targeted region both in sequence and configuration explain the serotype-specificity of M170. In addition, residues comprising the epitopes of M170 are highly conserved across currently circulating type O isolates, suggesting that M170 is likely capable of neutralizing most FMDV type O strains (Supplementary Fig. 8).

### Structural basis for the protective efficacy of M8 and M170 against FMDV infections

Competitive SPR and cell-based viral attachment inhibition assays demonstrated the abilities of M8 and M170 to effectively abrogate the interactions between FMDV and its receptor integrin (Fig. 3). In addition to integrin, FMDV can often readily adapt to tissue culture, where infection occurs in the absence of integrin via acquired basic mutations that confer an ability on the virus to interact with heparin sulfate (HS) (SaCarvalho et al., 1997). Cryo-EM structures of FMDV type O in complex with αvβ6, hitherto not at high resolution, reveal that the fully open form of the integrin attaches to an extended GH loop of VP1 via interactions with the RGD motif plus downstream hydrophobic residues and a conserved previously identified HS binding site through an N-linked sugar (Kotecha et al., 2017). Superposition of the FMDV-M8/FMDV-M170 and FMDV-αvβ6 complex structures revealed clashes between the two subunits of αvβ6 and M8/M170 (Fig. 6a). Notably, the structural overlay analysis reveals that M8 competes with β6 subunit for binding to the RGD motif in a similar interaction mode and M170 partially occupies the αv subunit binding site on the virus, where an N-linked sugar from the αv is oriented (Fig. 6a). Thereby, binding of M8 or M170 can completely block the attachment of integrin to FMDV owing to the substantially stronger binding affinities of M8 or M170 for FMDV when compared to that of the integrin. In addition, M170 targets the HS binding site, occluding the infection mediated by sulfated sugars as well (Fig. 6b). Many picornaviruses have been shown to harbor concealed binding regions for their receptors. For example, some enteroviruses, harness the “canyon” as their receptor-binding site, inside which the receptors have to insert (Kang Wang, 2020b; Ren et al., 2013). This prevents the recognition of the virus by most antibodies thereby assisting the virus in escaping the NAbs. In contrast to concealing receptor binding sites, some picornaviruses like FMDV and hepatitis A virus (HAV) expose their receptor binding sites and most of their binding residues are remarkably non-conserved across receptor-dependent viruses (Kotecha et al., 2017; Wang et al., 2015; Zhou et al., 2019). However, essential residues for receptor recognition have been verified to be highly conserved and efficient (Kang Wang, 2020b; Zhou et al., 2019), such as the RGD motif, essential for attachment of many viruses to cell surface. Therefore, these viruses may evolve by mutating residues in the non-essential binding region to abrogate neutralization by their antibodies without interfering with receptor recognition. Notably, M8 and M170 recognize the RGD motif and the HS binding site, respectively, having great potential to occlude viral mutational escape. Footprints of αvβ6, HS, M8 and M170 on the FMDV surface reveal overlapped patches with areas of ~2,800 Å^2^, ~1,900 Å^2^ and ~120 Å^2^, between αvβ6 and M8/ M170, and between HS and M170, respectively (Fig. 6c and 6d). Collectively, the abilities of M8 and M170 in preventing FMDV from attaching to host cell surface can be attributed to steric clashes arising out of partially overlapping binding sites.

**Fig. 6.**
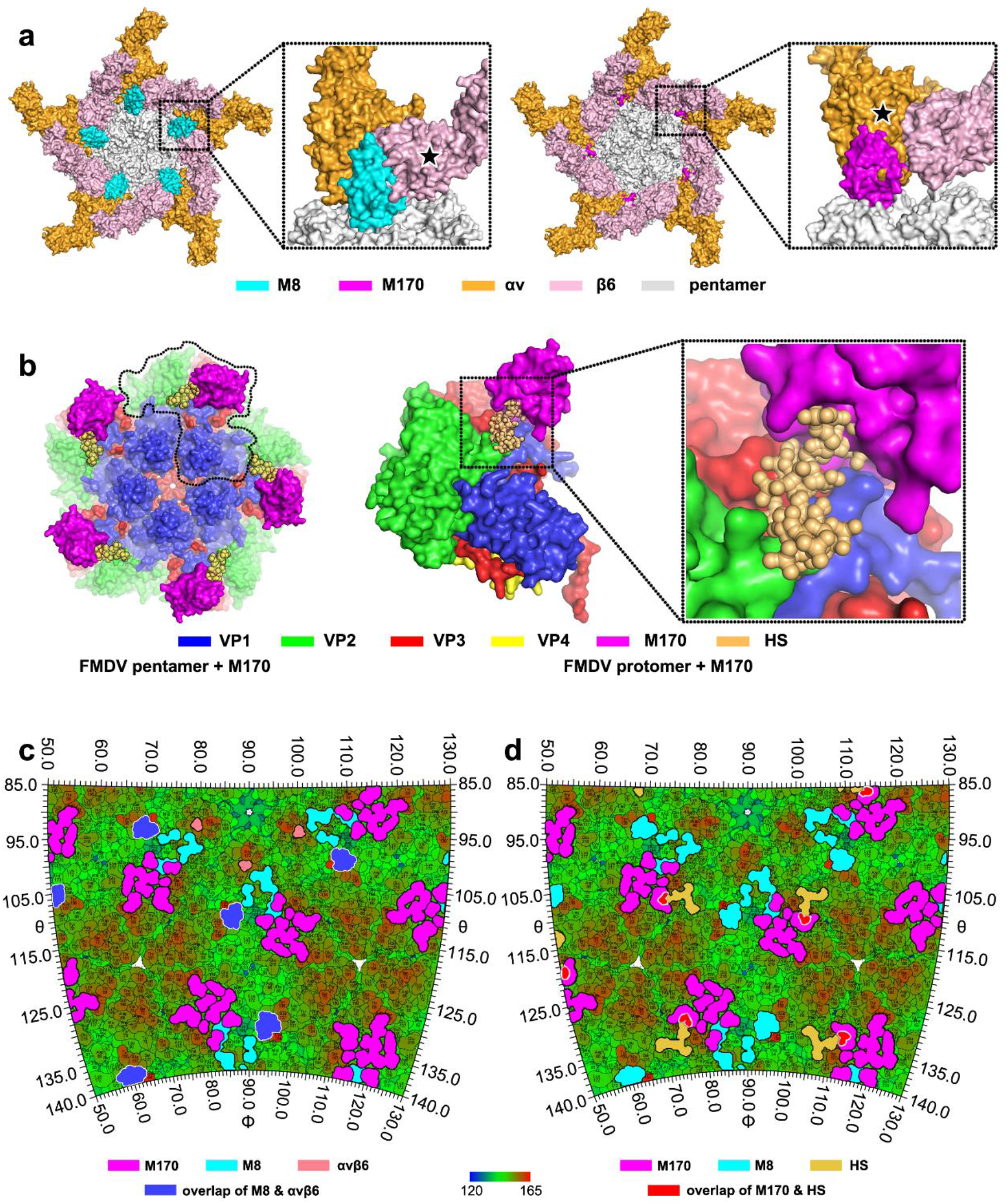
Mechanism of neutralization of FMDV by M8 or M170. **a** Clashes between M8 (left)/M170 (right) and αvβ6 receptor. Superimpositions of structures of the FMDV-M8/M170 and FMDV-αvβ6 are shown in pentamer. The capsid pentamer, M8, M170, and receptor subunits αv and β6 are colored in grey, cyan, magenta, bright orange and light pink, respectively. Clashes between M8/M170 and receptor subunits are prominent and marked with star symbols. **b** Clashes between M170 and HS cellular receptor. Color scheme for capsid pentamer and M170 is same as in Fig. 4d. HS receptor is shown as ball and colored in yellow. A protomer together with its corresponding binders (HS and M170) is marked by black lines. The inset shows the clashes between HS and M170. **c and d** Roadmap showing the relative positions of the NAb (M8 and M170) and receptor (αvβ6 and HS) footprints on the viral surface. The footprints of M8, M170, αvβ6 and HS receptor are outlined with cyan, magenta, light pink and yellow line, respectively. Overlapped footprints between M8 and αvβ6 and between M170 and HS are highlighted in blue and red.

## Discussion

FMD outbreaks are reported every year in countries spread across continents like Asia, Africa, South America and Europe. Vaccination is a major preventive strategy to control FMD and chemically inactivated FMDVs formulated with an adjuvant are routinely administered in populations at a high risk of contracting the disease. However, FMD vaccine stability is a big issue since intact virions with a sedimentation coefficient of 146S can irreversibly dissociate into stable pentamers, referred to as 12S viral capsid degradation products, leading to dramatically decreased immunogenic potency of these vaccines (Paton et al., 2009). It is therefore critical to quantify and continuously monitor the 146S particles present in the crude FMDV antigen preparations used for vaccine manufacturing. Several approaches have been established to measure the concentration of 146S particles present in a crude FMDV preparation, which include traditional sucrose density gradient ultracentrifugation, size-exclusion, high-performance liquid chromatography and recently developed antibody-based ELISA (Harmsen et al., 2017). Among these, double antibody sandwich ELISA has advantages of being more sensitive, amenable to being adopted in a high-throughput mode and with an ability to discriminate between various serotypes in multivalent vaccines (Harmsen et al., 2017). A prerequisite to the development of ELISA-based tests for the quality control of FMD vaccines is the availability of antibodies that can specifically detect 146S particles. Antibodies that specifically bind to 146S particles are rarely reported because most of the mAbs can bind to both 146S and 12S particles. In our previous study we reported that M8, like most mAbs, has no clear binding preference or specificity for 146S or 12S particles. However, M170 specifically interacts with 146S particles, and cannot bind 12S particles directly (Supplementary Fig. 9) (Harmsen et al., 2011). Atomic details of the FMDV antigenic sites for M8 and M170 revealed in this study provide structural evidences underpinning the binding preferences of these two antibodies. Residues of the virus involved in binding M8 cluster into a patch within the pentamer, between the fivefold and twofold axes, mainly involving the VP1 GH loop (Fig. 4). Distinct from M8, M170 binds to the viral surface adjacent to the edges of pentameric building blocks nearby the twofold axes, primarily associating with a number of exposed loops of VP3 (Fig. 4). During the de-assembly of the 146S particle into 12S pentamers, the capsid undergoes significant conformational rearrangements accompanied by a ~4.6° rotation of the bulk of VP3 towards the periphery, giving rise to a structural shift of some distal loops, including EF, HI, DE and BC loops, in the range of 4-9 Å (Supplementary Fig. 9). Remarkably and in line with the inability of M170 to recognize 12S particles, some residues of the M170 epitope, such as D71 and V73 in BC loop, E131 and K134 in EF loop, lie in the region with the largest conformational changes (Supplementary Fig. 9). In contrast to M170, M8 shows indistinguishable binding activities against 146S and 12S since its epitope is located beyond this region (Supplementary Fig. 9). Therefore, the 146S-specific M170, together with 12S-directed antibody, can be combined to develop an ELISA system for the quantification of 146S particles during vaccine manufacturing. The molecular features of the M8 and M170 epitopes unveiled in this study pose interesting targets for structure-based rational broad-spectrum vaccine design and effective antigen detection development for use in vaccine quality control. Our studies also highlight the promise of simultaneous administration of vaccines and therapeutic antibodies for rapid and complete protection prior to the establishment of effective immune responses elicited by FMDV vaccines.

## Materials and methods

### Virus production and purification

FMDV serotype O strain O/BY/CHA/2010 (GenBank accession no. JN998085.1) from the OIE/National Foot-and-Mouth Disease Reference Laboratory (Lanzhou, China), was cultured in Baby Hamster Kidney (BHK)-21 cells at 37°C for 8-10 h with Dulbecco’s modified Eagle’s medium (DMEM) (Gibco, CA, USA) supplemented with 100 U/ml penicillin, 100 μg/ml streptomycin, and 2% fetal bovine serum (FBS, Gibco). After removing the cell debris, the virus supernatant was concentrated using 8% polyethylene glycol (PEG)-6,000 (Sigma-Aldrich, U.S.A). The pellet was resuspended in TNE buffer (50 mM Tris-HCl, 1 mM EDTA, 150 mM NaCl, pH 7.6), and mixed with an equal volume of trichloromethane and then centrifuged at 100,000 g for 1 h. Rude virus solution was loaded onto a 15-45% (w/v) sucrose density gradient, ultra-centrifuged at 120,000 g for 3 h with Beckman SW41 rotor. Target fractions were collected and examined by negative-stain electron microscopy and SDS-PAGE.

### Production of M8 and M170 neutralizing antibodies (NAbs)

NAbs (M8 and M170) were isolated and sequenced from phage display immune libraries as previously described by M.M Harmsen (Harmsen et al., 2005). NAb encoding genes were inserted between the *BamH* I and *Xho* I sites of plasmid pGEX-4T-1, and expressed with an N-terminal GST tag in *Escherichia coli* BL21(DE3) cell. Recombinant proteins were purified by GSTrap HP column (5 ml) on AKTA Pure system (GE), according to the manufacturer’ s instructions. Thrombin (Sigma, USA) was used to cleave the GST tag and an extra step for GSTrap HP column purification was performed to remove cleaved GST tag.

### Binding abilities of NAbs to FMDV

The binding abilities of M8 and M170 to FMDV were measured by double antibody sandwich ELISA (Harmsen et al., 2005). Briefly, 96-well plates were coated with 50 ng antibodies (M8, M170, 0.5 μg/ml) per well in coating buffer solution overnight at 4 °C. The plates were filled with 50 μl of 3-fold serially diluted FMDV (initial concentration 3 μg/ml), including O1/BFS1860/UK/67, A TUR/20/2006, A24/Cruzeiro/BRA/55, Asia 1/Shamir/ISR/89, C1/Detmold/FRG/60, and incubated at 25 °C for 1 h. The plates were washed for three times and further incubated with biotinylated NAbs (0.25 μg/ml) at 25°C for 1 h. After washing plates, PO-conjugated streptavidin (Jackson Immunoresearch, USA, lot no 79940) was added for another 1 h at 25 °C. Then, the plates were stained with 3,3’,5,5’-tetramethylbenzidine (TMB), and absorbance was measured at 450 nm in a 96-well plate reader.

### Plaque-reduction neutralization assay

The plaque-forming unit (PFU) of the viable virus neutralized by NAbs (M8 and M170) was determined using plaque assay. Antibodies were 2-fold serially diluted with the highest concentrations of 51.2 μM, or the cocktail of M8 and M170 (One-third of the neutralizing titer of the antibody plus a 2-fold diluted another antibody), which were incubated with an equal volume of virus (200 PFU/ml) at 37 °C for 1 h. The complexes were transferred to a 6-well culture plate with the momolayer BHK21 cells. After further incubating the plate for 1 h with slightly shaking every 20 min, the survival virus attached to the cell surface. The plates were covered with the gum tragacanth (2 ml/well) supplemented with 2% FBS and further incubated for 72 h. After removing the overlay, the cells were washed with PBS and fixed with 4% paraformaldehyde. Plaques were visualized by staining with 2.5% crystal violet. According to the neutralization efficiency of the experimental group to the control sample, the neutralization titers of the two antibodies were calculated by nonlinear curve fitting. The Data is presented as the mean ± SD of triplicate measurements.

### Binding affinity measurements

SPR experiments were performed by a BIAcore 8k machine with CM5 sensor chips (both GE Healthcare) in PBST buffer (PBS, 0.05% Tween-20(v/v)). Due to the requirement of acidic condition for the sensor-labeling, NAbs or receptors were loaded onto the sensor. The serially diluted FMDV (0, 2.5, 5, 10, 20 and 40 nM) flowed over the NAb (M8 or M170)-immobilized CM5 sensor chip surface. The binding affinities of two NAbs were analyzed using the software BIAevaluation Version 4.1. To further analyze the ability of two NAbs simultaneously to bind the virus, inactivated FMDV viruses firstly flowed through one NAb-immobilized CM5 chip. Prior to the other NAb flowing through, the same NAb acted as the flow phase to fully occupy the binding sites on virus surface, and then the other NAb flowed through for binding signal detecting. For blocking virus-receptor interaction, αvβ6 integrin receptor (R&D, lot: DCHM0319011) was immobilized on CM5 chips at concentrations equivalent to ~250 response units. Mixtures of 20 nM FMDV and different concentrations of NAbs flowed over the chip. Binding signals were detected and analyzed with the software BIAevaluation Version 4.1.

### FMDV challenge in guinea pigs

Guinea pigs (weight 300-400 g) were randomly divided into six groups, including therapeutic and prophylactic groups. Briefly, groups of guinea pigs were administrated intramuscularly with M8/M170 (2.5 mg/kg) 1 day before (prophylactic) or after (therapeutic) challenge with 100 50% median infective doses (100 ID_50_) of FMDV (0.2 ml) on the left hind footpad. Guinea pigs injected intramuscularly with PBS before or after challenge were acted as control groups. All animals were examined for clinical symptoms at 1 to 10 days post-infection (DPI). No lesions were considered as full protection, and blood samples were collected for detecting the viral RNA copies by real-time quantitative PCR (RT-qPCR). In brief, the total RNA of samples was extracted by TRIzol reagent (Invitrogen) for the synthesis of cDNA using PrimeScript^™^ RT Master Mix (TaKaRa, Dalian, China). RT-qPCR was performed on a CFX96 Touch™ Real-Time PCR Detection System (Bio-Rad Laboratories, Hercules, CA, USA), by 40 cycles of denaturation at 95 °C for 30 s, annealing and extension at 60 °C for 30 s. A total of 20 μl reaction system contains 10 μl 2X Premix Ex Taq II (TaKaRa, Dalian, China), 1 μl of 3D gene specific primer (forward: 5’ ACTGGGTTTTACAAACCTGTG A 3’; reverse: 5’ GCGAGTCCTGCCACGGA 3’), 2 μl of fluorescent probe (5’ TCC TTT GCA CGC CGT GGG AC 3’), and 2 μl of the cDNA template. The pcDNA3.1-3D plasmids was constructed and quantified as a standard sample.

### Virus quantification on the cell surface by RT-PCR

The amounts of FMDV remaining on the surface of BHK21 cells after M8/M170 treatment were estimated using qPCR as previously described. Briefly, FMDV was incubated with the serially diluted NAbs before and after the virus attached to BHK21 cells (MOI=1) at 4 °C. The cells were washed three times and the total RNA was extracted by TRIzol reagent (Invitrogen). The cDNA was prepared by PrimeScript^™^ RT Master Mix (TaKaRa, Dalian, China). The level of virus mRNA was quantified using SYBR Premix Ex Tag II (Tli RnaseH Plus) on CFX96 Touch™ Real-Time PCR Detection System (Bio-Rad Laboratories, Hercules, CA, USA), the reaction system is the same as before. The level of glyceraldehyde-3-phosphate dehydrogenase (GAPDH forward: 5’ AAGAAGGTGGTGAAGCAGGCATC 3’, GAPDH reverse: 5’ CGCCATCGAAGGTGGAAGAGTG 3’) was used as an internal control. The relative levels of mRNA in different samples were represented using the 2^-ΔΔct^ method (Livak and Schmittgen, 2001).

### Thermofluor Assay

Thermofluor assay was performed with a MX3005p RT-PCR instrument (Agilent), SYTO9 (invitrogen) was used as fluorescent probe to detect the single-stranded RNA from virus capsid. In brief, the 50 μl reaction system includes, 2 μg purified viruses or 2 μg of viruses plus 1.5 μg of NAbs (~120 antibody molecules per FMDV virion), or 37 °C treated viruses or virus-antibody complexes and 5 μM SYTO9 in PBS buffer solution. System program was ramped from 25 to 99 °C with fluorescence recorded in triplicate at 1 °C intervals.

### Cryo-EM and data collection

Purified M8/M170 were incubated with purified FMDV particles (at a concentration of 0.5 mg/ml) at 4 °C for 1 min at the ratio of ~300 NAbs per FMDV particle. A 3 μl aliquot of the complex of FMDV and M8/M170 were applied to a freshly glow-discharged 400-mesh holey carbon-coated copper grid (C-flat, CF-2/1-2C, Protochips). Grids were blotted for 3 s in 90% relative humidity for plunge-freezing (Vitrobot; FEI) in liquid ethane. Cryo-EM datasets of FMDV-M8 and FMDV-M170 were collected with Talos Arctica and Titan Krios microscopes (FEI), both of which were equipped with a direct electron detector (K2 Summit; Gatan). Movies ((25 frames, each 0.2 s, total dose 30 e^-^ Å^-2^) were recorded with a defocus between 1.2 and 2.8 μm. Automated single-particle data acquisition was performed by SerialEM (Mastronarde, 2005), with a calibrated magnification of 59,000 yielding a final pixel size of 1.32 Å and 1.35 Å for FMDV-M8 and FMDV-M170, respectively.

### Image processing, model building and refinement

A total of 2,493 micrographs (FMDV-M8 complex) and 298 micrographs (FMDV-M170 complex) were recorded, respectively. Frames were corrected for beam-induced draft by aligning and averaging the individual frame of each movie using MOTIONCORR2 (Li et al., 2013). The contrast transfer function parameters were estimated by Gctf (Zhang, 2016). Particles were picked manually by Manual pick in RELION3 (Scheres, 2012). A total of 5,508 particles and 3,800 particles for FMDV-M8 and FMDV-M170 complexes were picked, respectively and selected to two-dimensional alignment and three-dimensional reconstruction. Finally, 3,701 particles of FDMV-M8 complex and 3,692 particles of FMDV-M170 complex were used for the icosahedral symmetry reconstruction. The resolution of the final icosahedral reconstructions was 3.2 Å and 3.1 Å, as evaluated by Fourier shell correction (threshold = 0.143 criterion). Although the overall resolution for these two icosahedral reconstructions is up to 3.1 Å – 3.2 Å, the maps for the binding interface between FMDV and NAbs are quite weak due to the relative low occupancy of NAbs and conformational heterogeneity. To improve the resolution for the binding interface, we used the block-based reconstruction strategy for focusing classification and refinement. The orientation parameters of each particle determined in Relion were used to guide extraction of the block region (~50% bigger than protomer-NAb) and these blocks were further 3D classified. A local reconstruction focusing on the protomer-NAb region was carried out, yielding a resolution of 3.9 Å and 3.5 Å for the interface of FMDV-M8 and FMDV-M170, respectively. The atomic model of FMDV (PDB code: 5DDJ) was initially fitted into our maps with CHIMERA (Pettersen et al., 2004) and future corrected manually by real-space refinement in COOT (Emsley and Cowtan, 2004). The atomic models of M8 and M170 were built *de novo* into densities with structures of single-domain antibodies as a guide, using COOT. These models were further refined by positional and B-factor refinement in real space with Phenix (Afonine et al., 2012). Refinement statistics are summarized in Table S1.

## Acknowledgements

We thank B. Zhu, X. Huang and G. Ji for Cryo-EM data collection at the Center for Biological imaging (CBI), Institute of Biophysics, and Y Chen, Z. Yang and B. Zhou for SPR technical support and guidance. We also thank P. Du, X.Y Zhi, Y Zhang for their guidance in preparation and purification of virus and antibody, and S.H Yin, J.X Ru for their help and guidance in guinea pigs experiments.

## Author contributions

H.D., P.L., and M.B. performed the experiments; P.L. and K.W. collected Cryo-EM data; P.L. and X.W. solved the structure; M.M. provided suggestion and some data about NAbs; D.Z. and R.F. did SPR experiments; S.M. and H.L help for doing some biological function verification experiment, S.S., X.W., and H.G. designed the study, H.D., S.S., and X.W. analyzed data, and H.D., X.W. wrote the manuscript, M.M., S.S., and H.G revised manuscript.

## Data and materials availability

The cryo-EM density maps of icosahedral reconstructions for FMDV-M8 and FMDV-M170 and blocked reconstructions for the binding interface of FMDV-M8 and FMDV-M170 have been deposited in the Electron Microscopy Data Bank under accession codes: EMD-WWWW, EMD-XXXX, EMD-YYYY and EMD-ZZZZ, respectively. The corresponding atomic coordinates have been submitted to the Protein Data Bank with accession numbers: 7WWWW, 7XXX, 7YYY and 7ZZZ, respectively.

## Abbreviations

FMDV: foot-and-mouth disease virus
PEG: polyethylene glycol
PFU: plaque-forming unit
DPI: days post-infection
100 ID50: 100 50% median infective doses
MOI: multiplicity of infection
SPR: surface plasmon resonance
Cryo-EM: cryo-electron microscopy
NAbs: neutralizing antibodies
RGD: Arg-Gly-Asp
ELISA: enzyme-linked immunosorbent assay
Neut50: 50% neutralizing concentration value
CDR: complementary determining regions
GP: guinea pigs
EVD68: Enterovirus D68
E30: Echovirus 30
EV71: enterovirus A71
RT-PCR: real-time reverse transcriptase–PCR
HS: heparin sulfate
P: prophylactic
T: therapeutic
PI: post infection

## Compliance with Ethics Guidelines

Hu Dong, Pan Liu, Manyuan Bai, Kang Wang, Rui Feng, Dandan Zhu, Yao Sun, Suyu Mu, Haozhou Li, Michiel Harmsen, Shiqi Sun, Xiangxi Wang and Huichen Guo declare that they have no conflict of interest.

Animal experiments were carried out in accordance with the regulations for the administration of affairs concerning experimental animals approved by the State Science and Technology Commission of the People’s Republic of China and by the Committee for Animal Welfare and Safety in the Lanzhou Veterinary Research Institute of the Chinese Academy of Agricultural Sciences (No. LVRIAEC2018–008).

