## supplementary materials for "Structural and molecular basis for foot-and-mouth disease virus neutralization by two potent protective antibodies"

This file contains Supplementary Figures 1-9 and Supplementary Tables 1-3.

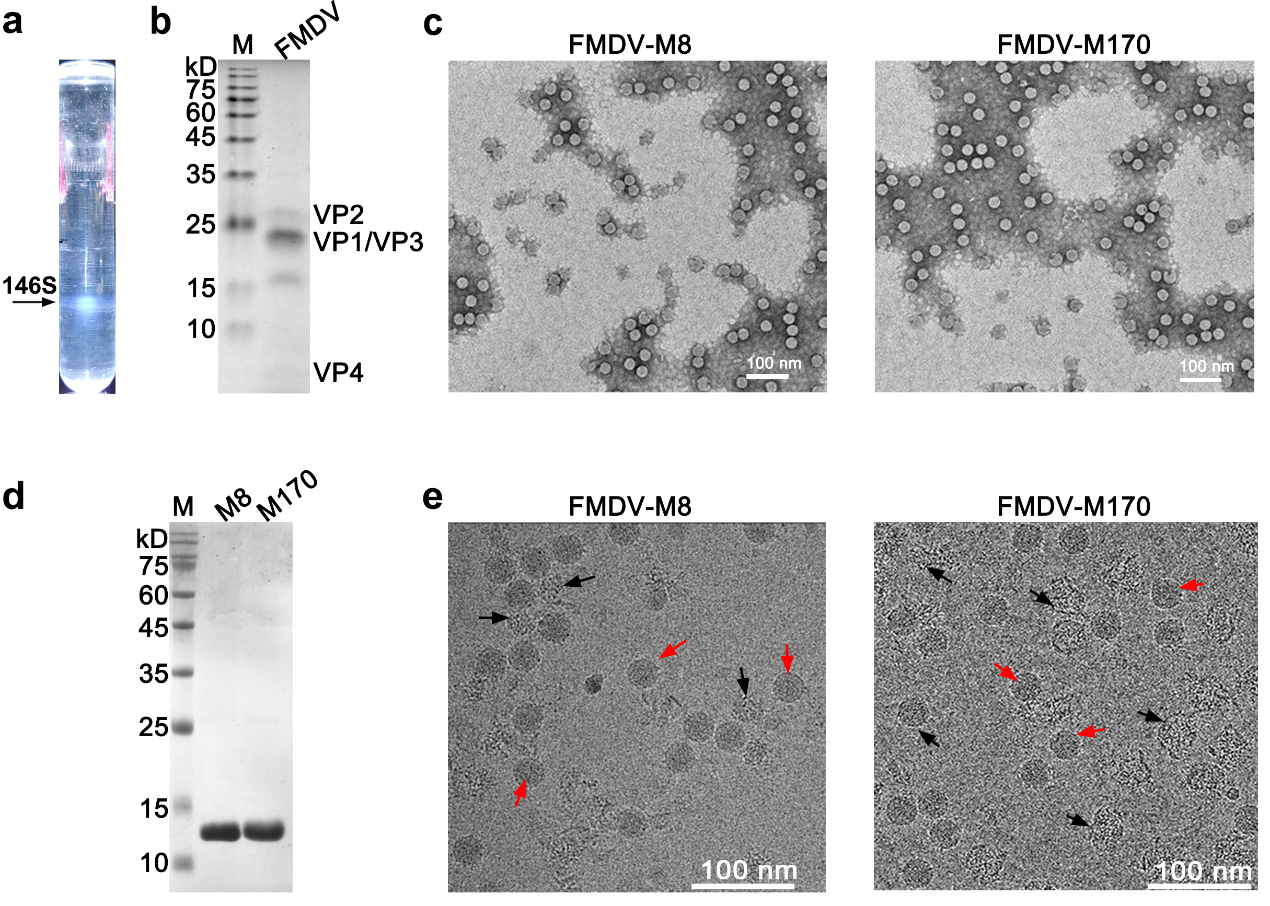

Supplementary Figure 1

**Purification and characterization of M8, M170 and FMDV.**

**a** Zonal ultracentrifugation of a 15 to 45% (w/v) sucrose density gradient at 120,000 g for 3 h was used to purify FMDV from the harvest concentrate described in the method section. Only one type of FMDV particle, corresponding to the 146S mature virion, was separated. **b** SDS-PAGE analysis for FMDV capsid proteins, the theoretical molecular weights of VP1, VP2, VP3 and VP4 are 23.7 kDa, 24.4 kDa, 23.9 kDa and 8.9 kDa, respectively. **c** The negative-stain images of FMDV in complex with M8 and FMDV in complex with M170. **d** Purity evaluation of M8 and M170 by SDS-PAGE. **e** The cryo-EM micrographs of FMDV-M8 and FMDV-M170 complexes, and the intact particles and broken particles were marked by red and black arrows, respectively.

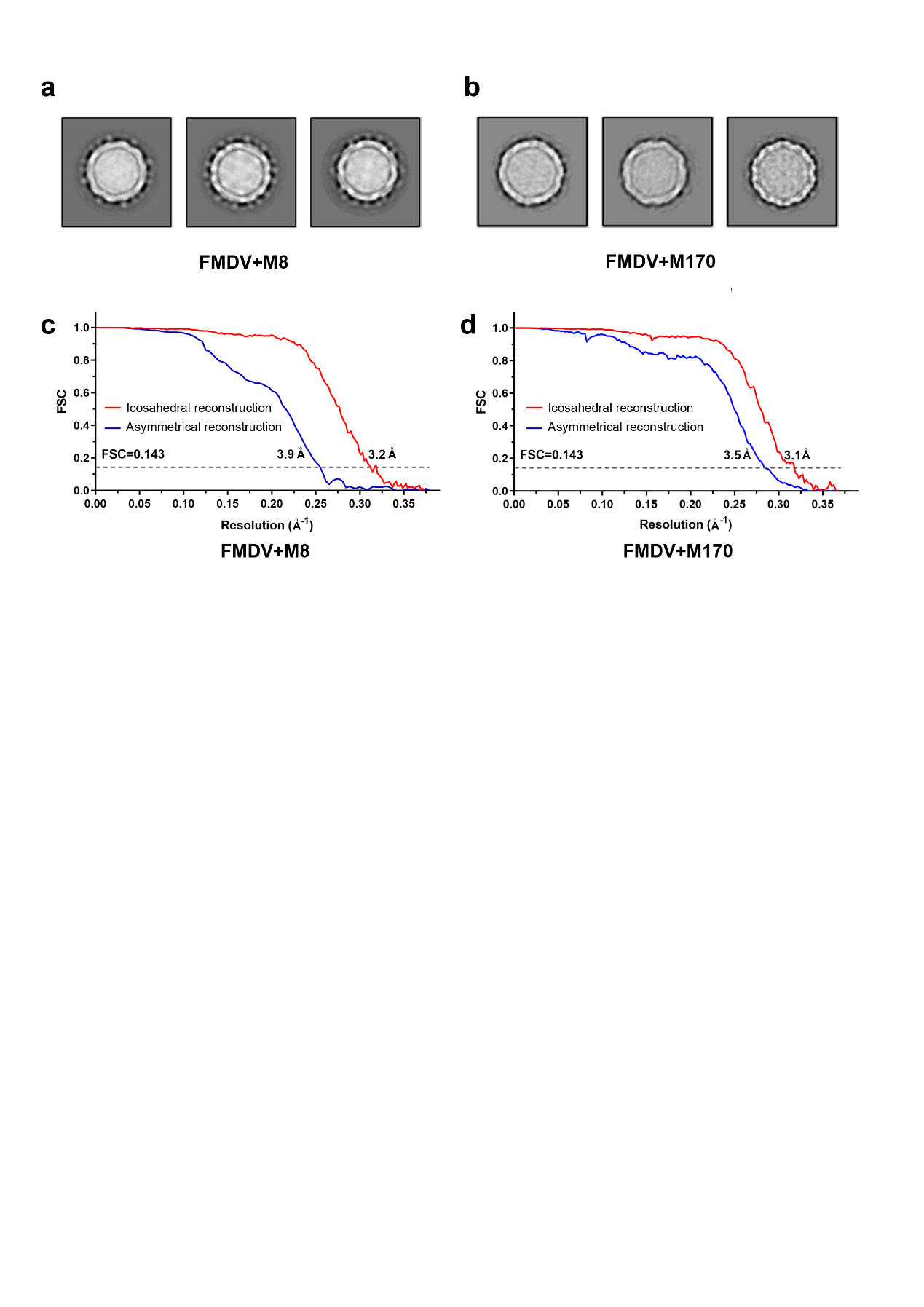

Supplementary Figure 2

**2D classification and FSC curves**

Representative classes from 2D classification in RELION for FMDV-M8 (**a**) and FMDV-M170 complexes (**b**). Gold-standard Fourier shell correlation (FSC) curves of the final maps of FMDV-M8 (**c**) and FMDV-M170 complexes (**d**).

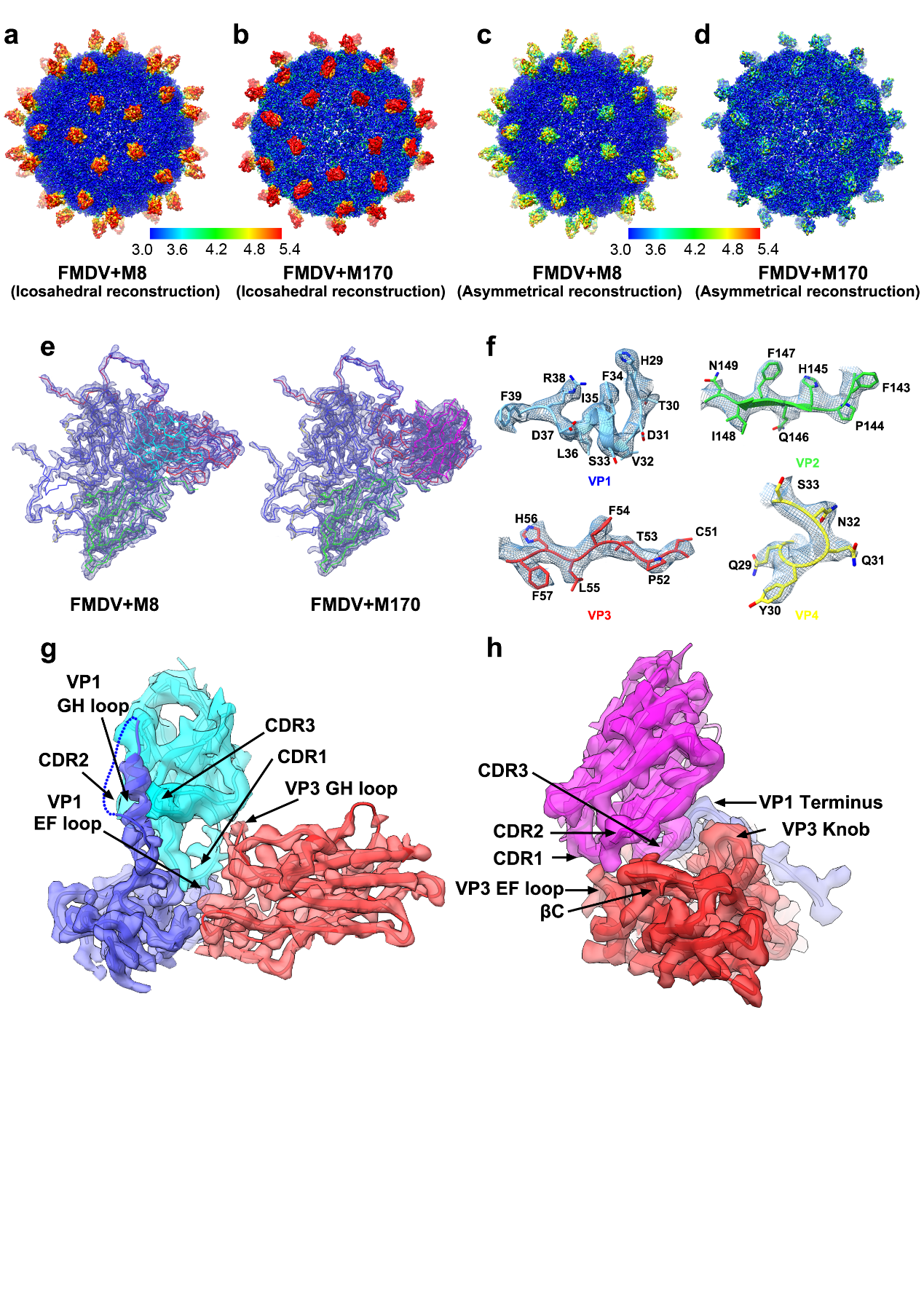

Supplementary Figure 3

**Map resolution evaluation and electron density maps**

Map resolution assessment of the icosahedral reconstructions of FMDV-M8 (**a**) and FMDV-M170 (**b**), and the asymmetrical reconstructions of FMDV-M8 (**c**) and FMDV-M170 (**d**) with color indicated below. Electron density maps for the FMDV protomer (**e**), the sidechains of VP1-VP4 (**f**) and the binding interface of M8 (**g**)/M170 (**h**) are shown.

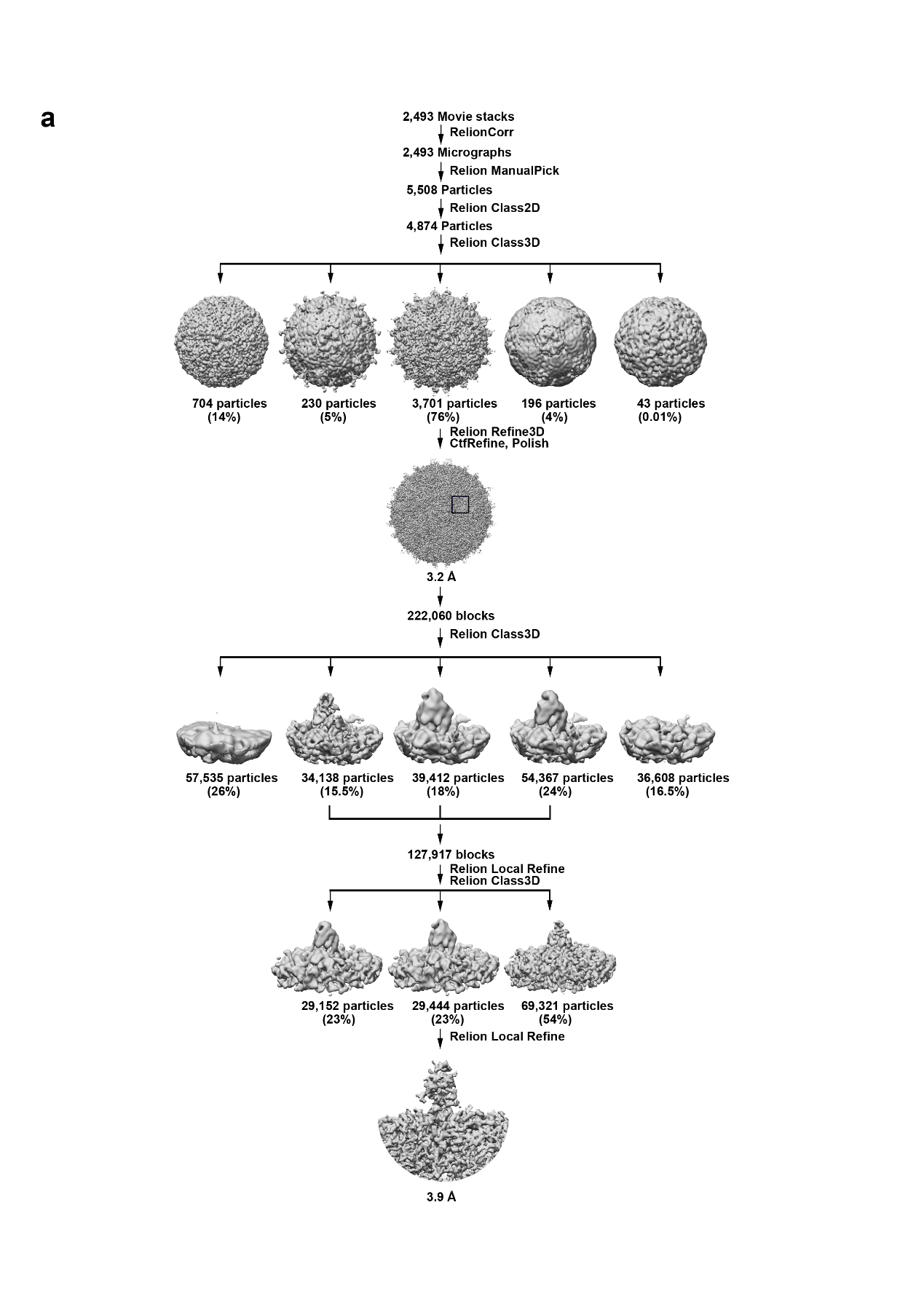

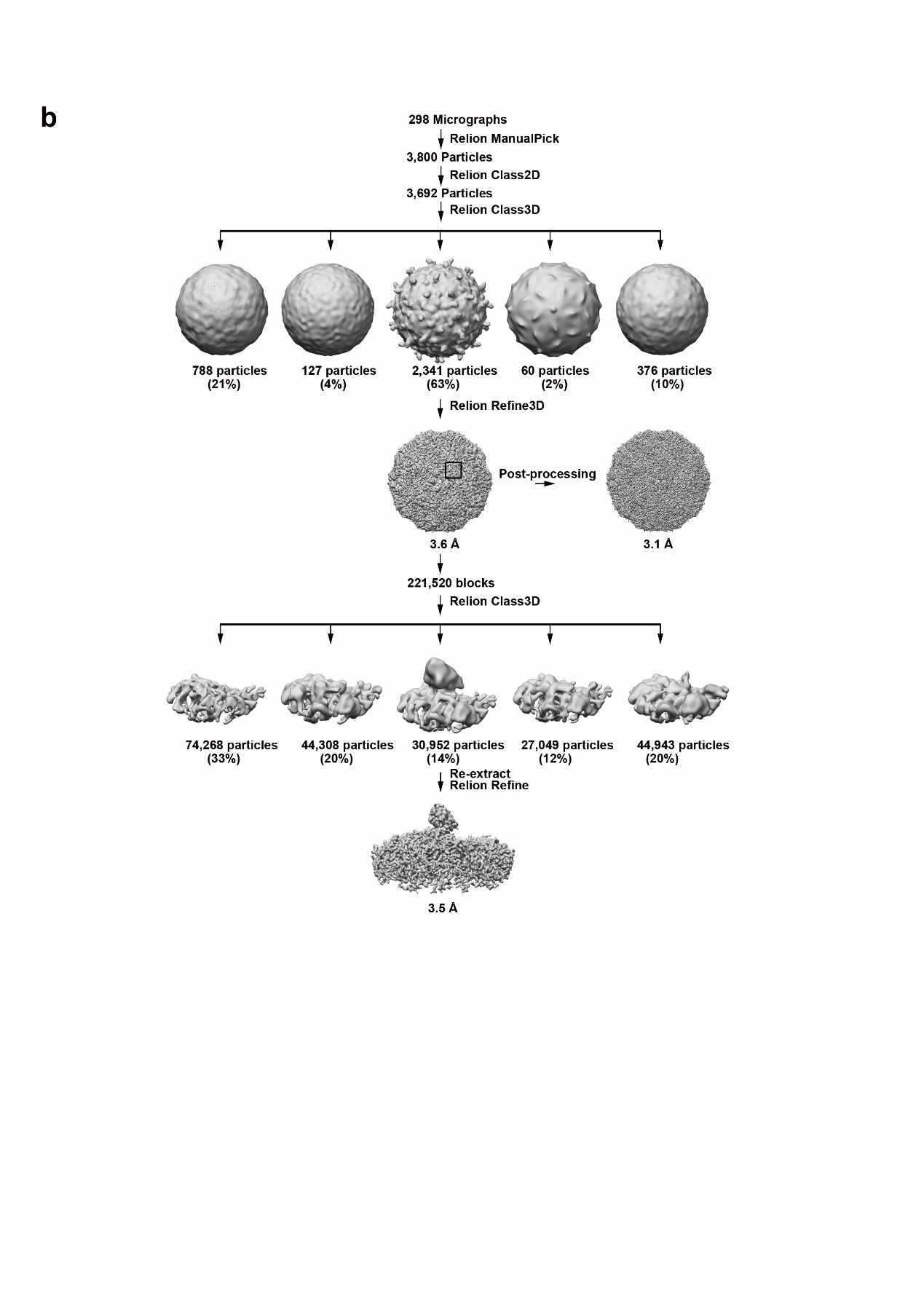

Supplementary Figure 4

**Flow-chart for Cryo-EM data processing**

(**a**) and (**b**) show the data processing procedures for FMDV-M8 and FMDV-M170 complexes, respectively. Details can be found in the Methods section.

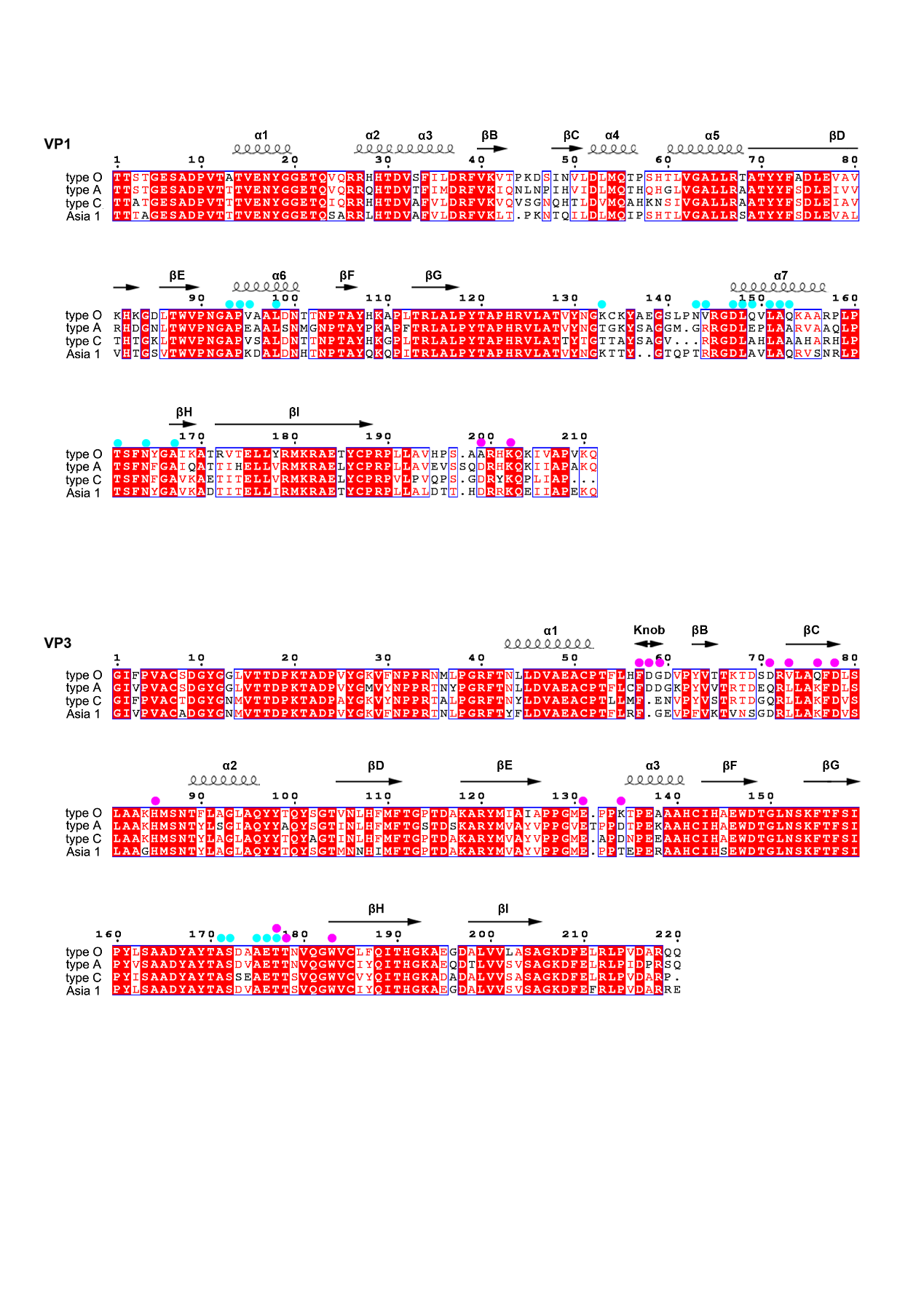

Supplementary Figure 5

**Sequence alignment of capsids between the four representative FMDV serotypes**

Sequence alignment of VP1 and VP3 from FMDV O with counterparts from 3 representative FMDV serotypes (A, C and Asia I). The residues involved in directly interacting with M8 and M170 are marked with cyan and magenta balls, respectively.

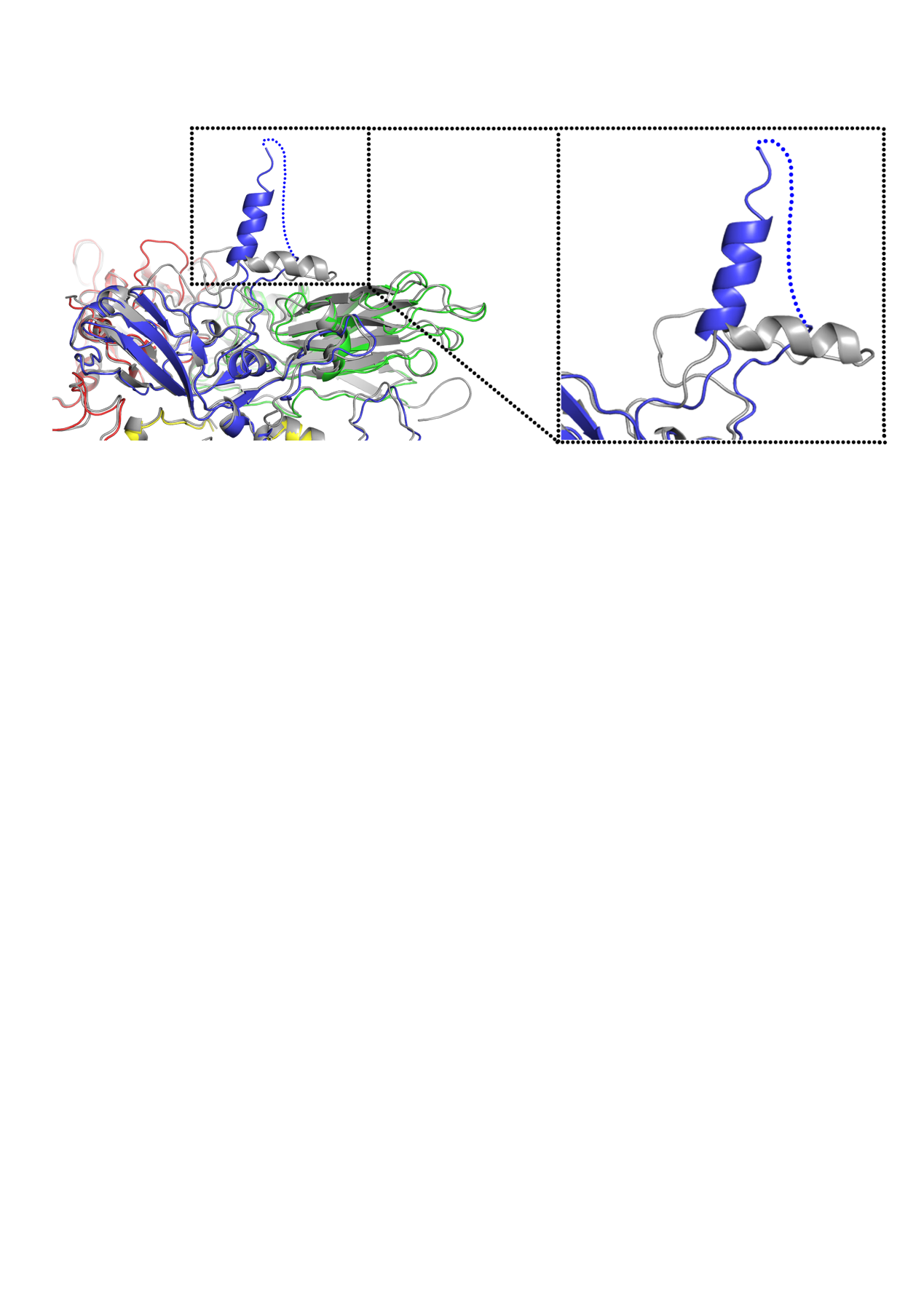

Supplementary Figure 6

**Conformational comparison of the VP1 GH loop**

The VP1 GH loop in the structure of FMDV-M8 complex exhibits an “up” configuration compared to the “down” conformation in the structure of the reductant treated FMDV (PDB Code: 1FOD). The color scheme for VP1-VP4 in the structure of FMDV-M8 complex is same as the Fig. 4b and the protomer from the reductant treated FMDV is colored in gray.

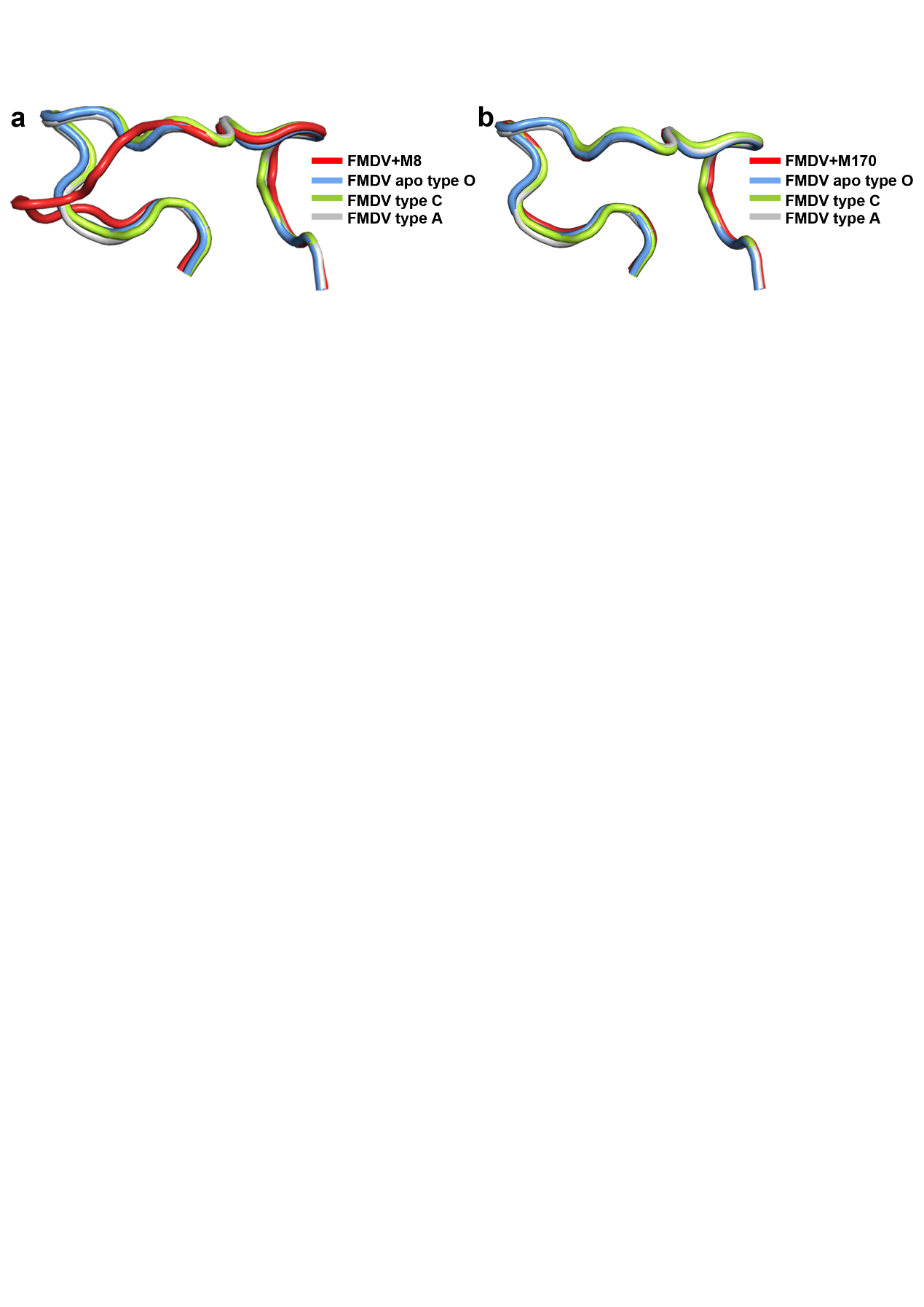

Supplementary Figure 7

**Conformation conservation analysis of the VP3 GH loop**

Structures of the VP3 GH loop from FMDV-M8 (**a**) and FMDV-M170 (**b**) are superposed with counterparts from 3 representative FMDV serotypes.

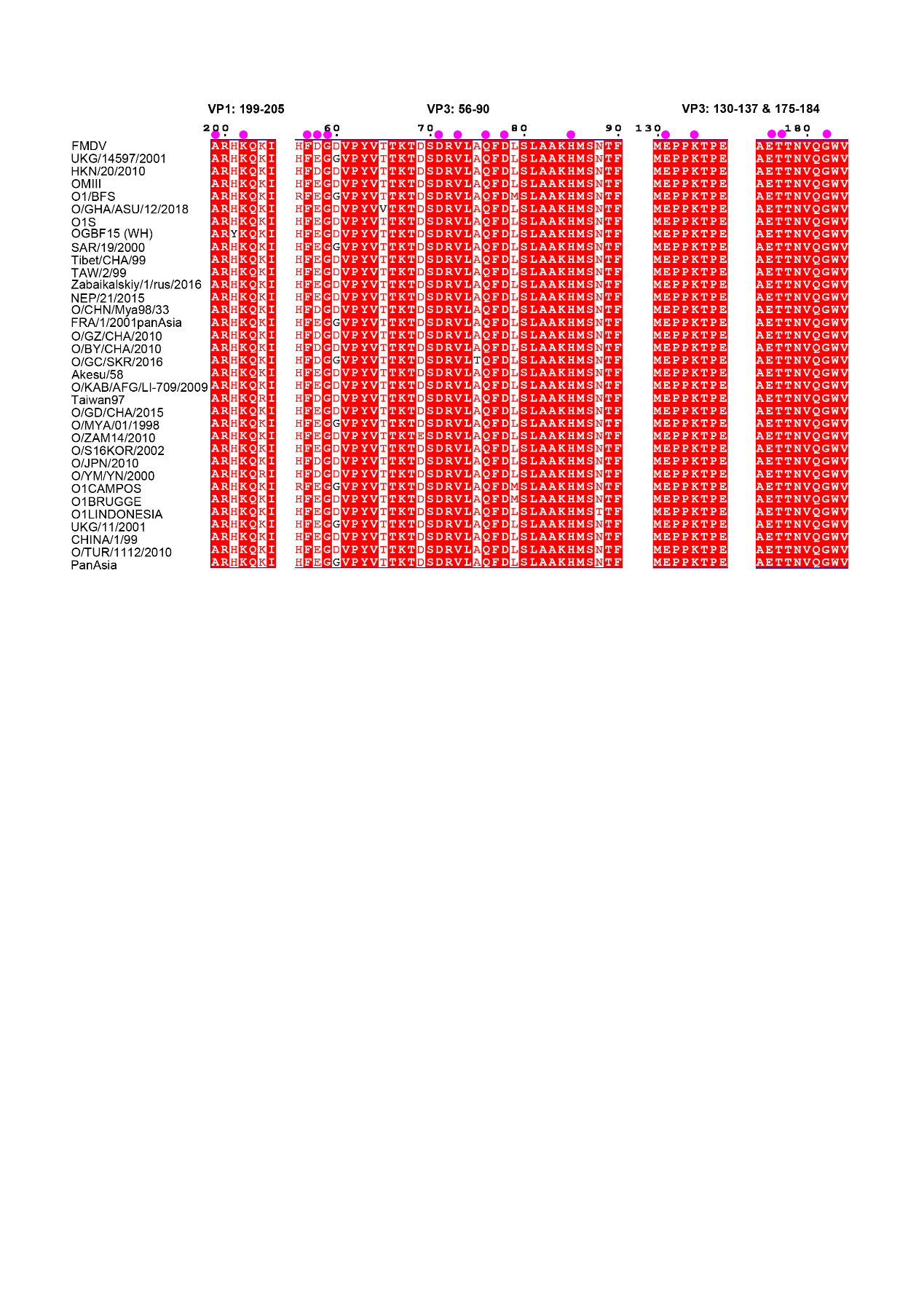

Supplementary Figure 8

**Multiple-sequence alignment analysis of the M170 epitope on FMDV O**

The alignment results are displayed with the program Espript.

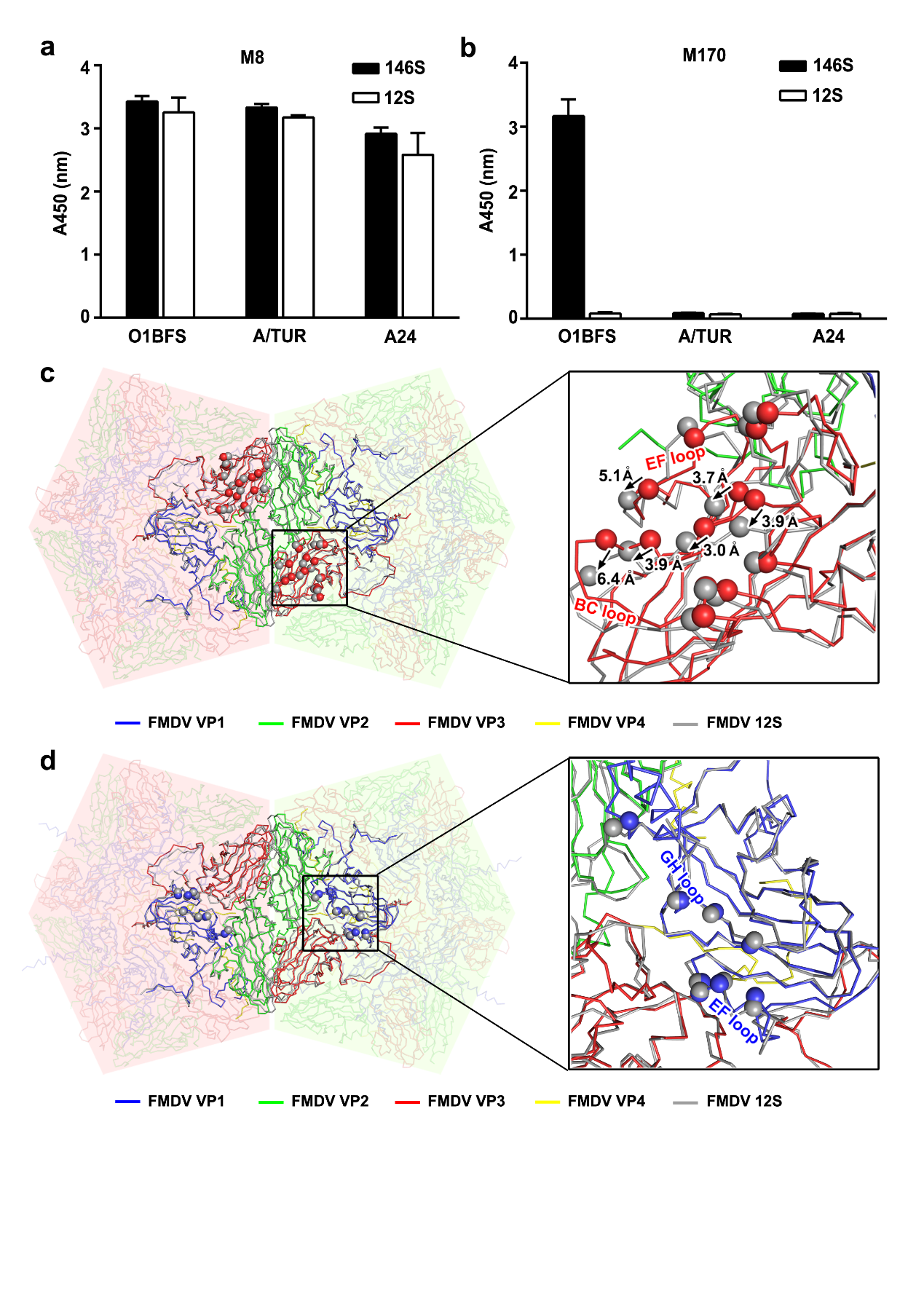

Supplementary Figure 9

**Structural basis of binding preference/specificity of M8/M170 for 146S or 12S particles**

**a** M8 exhibited no clear binding preference for 146S or 12S particles. **b** M170 showed specifically binding to 146S particles. Purified viruses (146S) and acid-dissociated pentamers (12S) from FMDV types A and O were used for testing the binding to M8 and M170 by ELISA. Structural analysis for explaining the binding specificity of M170 for 146S particles (**c**) and no binding preference of M8 for 146S or 12S particles (**d**). Superimposition of structures of the pentamers from 146S and 12S (PDB:5OYI) shows structural rearrangement in VP3, in particular the distal loops, such as BC and EF loops. Color scheme for the pentamer from 146S is same as Fig.4b and the structures of acid-dissociated pentamers are colored in gray. Epitopes of M8 and M170 are represented as spheres and colored corresponding to the protein chain they belong to. Major structural shifts are labeled and marked by black arrows.

| **Supplementary Table 1 Cryo-EM data collection and atomic models reﬁnement statistics** | | |
| --- | --- | --- |
| Data collection | | |
| Complex | FMDV+M8 | FMDV+M170 |
| Microscope | FEI Talos Arctica | FEI Titan Krios |
| Camera | Gatan K2 | Gatan K2 |
| Voltage (kV) | 200 | 300 |
| Total dose (e^-^/A^2^) | 30 | 30 |
| Micrographs (total) | 2,493 | 298 |
| Micrographs (used) | 2,493 | 298 |
| Particles selected | 5,508 | 3,800 |
| Particles included in final reconstruction | 3,701 | 3,692 |
| sampling, Å per pixel | 1.32 | 1.35 |
| Defocus range (μm) | 1.2-2.8 | 1.2-2.8 |
| Symmetry | I | I |
| Resolution (Å) (FSC=0.143 criterion) | 3.2 | 3.1 |
| Block particles included   in final reconstruction | 29,444 | 30,952 |
| Symmetry imposed on block particles | C1 | C1 |
| Resolution (Å) (C1 reconstruction) | 3.9 | 3.5 |
| Model refinement | | |
| Ramachandran statistics (%) | | |
| Most favored | 92.88 | 96.07 |
| Allowed | 6.36 | 3.93 |
| Outliers | 0.76 | 0 |
| R.m.s.d | | |
| Bond lengths (Å) | 0.014 | 0.008 |
| Bond angles (°) | 1.228 | 0.757 |

| **Supplementary Table 2. Residues of M8 interacting with the FMDV (d < 4 Å)** | | | |
| --- | --- | --- | --- |
| **FMDV** | | | **M8** |
| **Location** | **Domain** | **Residues** | **Residues** |
| **VP1** | **EF loop** | A93 | I34 N35 |
|  |  | P94 | S33 I34 |
|  |  | V95 | F32 S33 I34 N35 |
|  |  | L98 | N35 |
|  | **GH loop** | K133 | N110 |
|  |  | N143 | D69 S70 |
|  |  | V144 | Y67 D69 |
|  |  | D147 | A66 |
|  |  | L148 | A66 Y67 W113 |
|  |  | Q149 | A111 |
|  |  | L151 | T60 A66 W113 |
|  |  | A152 | N110 A111 W113 |
|  |  | Q153 | A111 |
|  |  | T161 | D36 I39 |
|  |  | N164 | D36 |
|  |  | A167 | N35 |
| **VP3** | **GH loop** | A171 | I34 |
|  |  | S172 | S33 I34 |
|  |  | A175 | G118 T119 |
|  |  | E176 | F117 |
|  |  | T177 | F117 |

| **Supplementary Table 3. Residues of M170 interacting with the FMDV (d < 4 Å)** | | | |
| --- | --- | --- | --- |
| **FMDV** | | | **M170** |
| **Location** | **Domain** | **Residues** | **Residues** |
| **VP1** | **C-terminus** | A199 | A104 P106 S108 |
|  |  | K202 | S108 |
| **VP3** | **Knob** | F57 | L105 |
|  |  | D58 | R53 L105 |
|  |  | G59 | Y62 |
|  | **BC loop** | D71 | W56 |
|  |  | V73 | W56 |
|  | **βC** | Q76 | A104 L105 |
|  | **CD loop** | D78 | F103 |
|  |  | H85 | L105 |
|  | **EF loop** | E131 | R30 S34 Y35 |
|  |  | K134 | W56 |
|  | **GH loop** | T177 | D109 Y110 |
|  |  | T178 | Y110 |
|  |  | W183 | F103 |
